# MARINER: a surround visual stimulator for vision research in aquatic animals

**DOI:** 10.64898/2026.06.09.731119

**Authors:** Tim C. Hladnik, David-Samuel Burkhardt, Yue Zhang, Patrick Weygoldt, Alexander Wendt, Buket Solak, Tod R. Thiele, Aristides B. Arrenberg

## Abstract

Aquatic vertebrates are increasingly used in neuroscience research, yet underwater visual stimulation remains a challenge. Commonly used monochromatic stimuli have been shown to be inadequate to activate many visual neurons properly, and underwater refraction artifacts are prone to ruin stimulus designs. Here, we present MARINER – a visual stimulator, which remedies these issues and integrates concurrent behavioral and neurophysiological two-photon calcium imaging recordings. MARINER’s full-field visual stimulation combined with receptive field mapping reveals that the visual field of larval zebrafish is larger than previously thought, extending almost down below the fish, and is spatially biased to better utilize motion content in naturalistic visual scenes. Using chromatic motion nulling, we further show that behavioral responses and task-associated sensory neurons are colorblind for “red” and “green” during the larva’s optokinetic response. The MARINER stimulator facilitates naturalistic stimulation and faithful presentation of colored visual underwater stimuli for small aquatic species.

## Introduction

Larval zebrafish have matured as a vision neuroscience model system in recent years. Their easy accessibility to neurophysiological imaging techniques enables researchers to study large neuronal populations of visuomotor circuits in live and behaving animals (Ahrens et al., 2013; Bahl & Engert, 2020; Brustein et al., 2003; Renninger & Orger, 2013; C. Yang et al., 2024; E. Yang et al., 2022). The easy optical access also fosters comparative studies on other aquatic species (Fouke et al., 2025; Mearns et al., 2025) and the phylogenetic position of fish provides a window into the circuit organization of a more ancestral vertebrate visual system with four cone photoreceptors (Baden & Osorio, 2019; Fornetto et al., 2025). However, accurate conclusions about vision circuits require precise knowledge on stimulation conditions during experiments to faithfully estimate sensory input. Much of previous research has focused on terrestrial species (Baden, 2020), where visual stimuli are comparatively easy to apply. Underwater visual stimulation, on the other hand, poses some critical challenges.

First, the interaction of light and the optical interfaces between different physical media causes refraction and reflection, which affects most aquatic laboratory settings, where light needs to travel through air, glass/plastic and water before reaching the animal’s eye. Second, ordinary LCD displays or video projection screens are designed for operation in air and cannot easily be used underwater. And third, the containers used to separate water from display, usually have at least one flat surface (e.g. the water surface), which can cause severe optical artifacts that jeopardize the visual stimulus design (Dunn & Fitzgerald, 2020; Wang et al., 2021). Taken together, these issues strongly weaken the reliability and interpretability of underwater vision experiments in potentially all aquatic species.

To add to this, many aquatic animals have laterally placed eyes, which makes development of stimulators that cover the whole (binocular) visual field challenging. Larval zebrafish have a large visual field of about 163° per eye (Easter, Jr. & Nicola, 1996), with only little binocular overlap. In effect, to study the fish’s whole visual field requires a stimulation system to provide almost complete 360° coverage. Gaining experimental access to the complete visual field is necessary, because visual function is highly anisotropic across the fish’s visual field. This anisotropy has been shown at the level of retinal structure (Yoshimatsu et al., 2020; Zhou et al., 2020), neuronal activity and circuitry (Förster et al., 2020; Wang et al., 2020; Zhang et al., 2022), as well as in innate behaviors (Bianco et al., 2011; Dehmelt et al., 2021; Kist & Portugues, 2019; Wang et al., 2020), and it is likely a result of adaptations to its habitat (Alexander et al., 2022; Zimmermann et al., 2018). Thus, any study of a particular circuit or behavior requires experimenters to be able to precisely stimulate the appropriate parts of the animal’s visual field.

Aquatic vertebrate species, such as amphibians and different taxa of fish, have different cone photoreceptor repertoires (Baden & Osorio, 2019; Loew & Lythgoe, 1978; Röhlich & Szél, 2000). Zebrafish vision, for example, is tetrachromatic (Branchek & Bremiller, 1984) with S and M-cone peak sensitivities that are shifted to somewhat shorter wavelengths compared to humans (Baden & Osorio, 2019; Endeman et al., 2013; Stockman et al., 2000). As a result, commonly available display devices aren’t suited to accurately present color stimuli to zebrafish, since they are optimized for human trichromatic vision (Franke et al., 2019). Recent work suggests (Fornetto et al., 2025), that completely disregarding chromatic contributions to vision may even be disadvantageous for parts of vision research that aren’t typically concerned with color processing, as it shows that monochromatic stimuli are less effective at driving motion responses, than polychromatic “white” ones.

To solve these challenges, we created MARINER – a Multispectral Arena for Real-time Interactive Neuronal and Ethological Research. It is a spherical visual stimulation arena which utilizes a single DLP video projector and an external, configurable light source to project images onto the outside coating of a transparent, spherical, and water-filled container. The projection covers 360° azimuth and 135° elevation and allows for simultaneous visual stimulation with three zebrafish cone-optimized spectra at a time. An openly available Python-/OpenGL software suite facilitates the mapping of arbitrary 3D visual stimuli onto the spherical surface and integrates visual stimulation with online tracking-based stimulus updating.

Here, we demonstrate the advantages of our novel stimulation arena by gaining insights into a range of hitherto largely inaccessible topics in zebrafish vision research. We start by exploiting the full-field stimulation area of our arena to map the spatial extents and local direction selectivity of receptive fields of motion-sensitive cells in the optic tectum and pretectum of larval zebrafish and end by employing multichromatic motion stimuli to investigate the contributions of different wavelengths to the optokinetic response and to the motion-sensitive neuronal circuits that drive it.

## Results

### Spherical arena enables full-field visual stimulation during behavioral and neuronal imaging

The MARINER setup is centered around the focal imaging volume of the microscope objective. The animal is mounted in a spherical, water-filled “fishbowl” (Figure 1A6 and Figure 1D). A single DMD video projector is used for visual stimulation (Figure 1A1 and B1). After the video projector, the image is split into four sub-images using a pyramid mirror and then reunited on the fishbowl to evenly stimulate the animal’s large visual field. The fishbowl’s outer surface is coated white and serves as a back-projection screen for the visual stimulus to avoid refraction and reflection artifacts that would otherwise occur at flat surfaces of a cuboid container. For accurate chromatic stimulation, the video projector is connected through a liquid light guide to a multispectral LED light source, the peak wavelengths of which were optimized for zebrafish cone isolation (S-Figure 1B).

**Figure 1.**
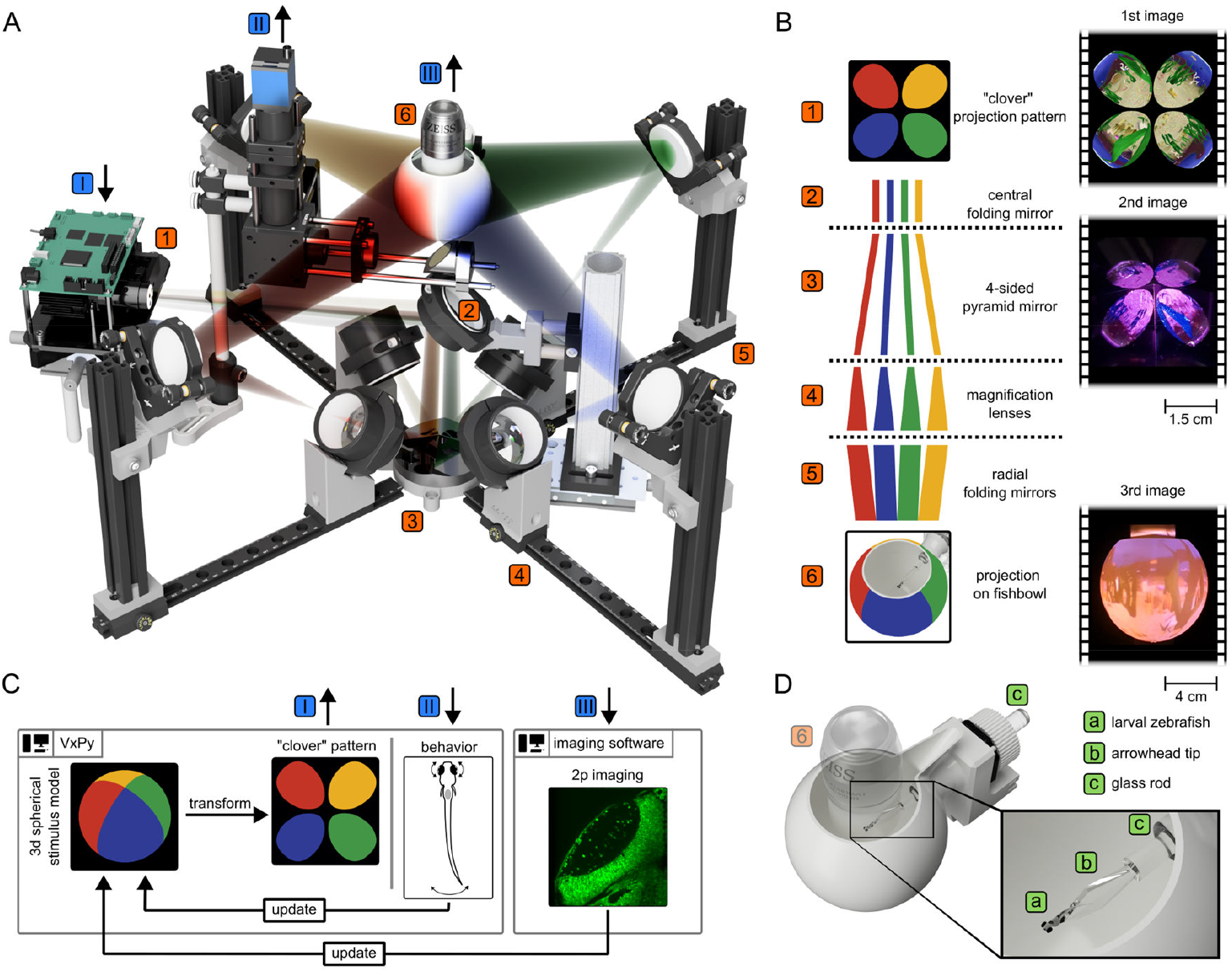
Overview of MARINER, a spherical visual stimulation arena. **A**: Bird view of the arena. Blue numerals mark inputs and outputs - (i) video projector for visual stimulation, (ii) camera for behavioral recordings and (iii) water-immersion objective for 2-photon imaging. Numbers 1 to 6 mark steps in the image transformation process in (A) and (B). **B**: Schematic of image transformation steps. The left side illustrates how the 4 different parts of the “clover” projection come together to form a seamless image on the spherical enclosure (“fishbowl”). The right side shows an example projection of a simulated VR underwater environment. The horizontally projected image (1) is reflected downward at a centrally located 45° folding mirror (2) and split into four sub-images by a pyramidally shaped mirror with four equal, 22° sloped sides at the bottom center of the setup assembly (3). Each of the four, radially outwards projected sub-images are then re-focused and magnified through a 2” achromatic lens (4) before being reflected at a78° folding mirror (5) towards the top center of the assembly, where the fishbowl with the embedded larva is located (6). **C**: Schematic of software components. Visual stimulus generation and behavioral recording are done in the custom-written Python package VxPy (Hladnik, 2025). 2-photon calcium imaging was done using the microscope vendor’s software (Sutter Instruments, MScan). **D**: Closeup of the fishbowl (number 6 in panel A, S-Figure 1A) with a zebrafish larva embedded in agarose (a) mounted on an arrowhead-shaped plastic stage (b) that is slotted into the conical cavity of a glass (or plastic) rod (c). The rod can be slotted through the rubber seal at the back of the fishbowl. This ensures that the animal is perfectly centered within the fishbowl and has an unobstructed 360° view of the visual scene.

To avoid light contamination of calcium imaging by the visual stimulation, we used a microcontroller circuit to turn the LEDs off in synchrony with the scanline periods or with the imaging frame rate, when using Galvo-Galvo or Resonant-Galvo scanners, respectively (S-Figure 1D). A small, round glass window at the bottom of the fishbowl, enables video recording and tracking of animal behaviors through an optical camera path from below (Figure 1A and C II, S-Figure 1C). Trans-illumination of the fish is achieved using an infrared light source (850nm) through the microscope’s objective.

Visual stimuli are generated in real-time based on a 3D spherical stimulus model, using a custom-written Python/OpenGL package (Hladnik, 2025). Stimuli may be directly constructed based on their location in a fish-centric unit sphere coordinate system, allowing for precise control over stimulus positions, orientations, and angular velocities (S-Video 2, CMN or moving grating stimuli). Alternatively, they may be created based on textures from secondary sources, such as, for example, footage from 360° cameras (S-Video 2, natural scenes), or cube map textures, thus enabling the rendering of simulated 3D virtual environments directly onto the spherical surface (Figure 1B, right column; S-Video 2 3D virtual reality).

### The zebrafish monocular visual field is about 180° in azimuth and almost reaches the nadir in negative elevation

Previous estimates of visual field size of larval zebrafish were based on geometrical considerations of eye and lens shapes (Easter, Jr. & Nicola, 1996) and therefore only coarse approximations. Our setup allowed measurement of the functional visual field by presenting stimuli at a high spatial resolution across almost their complete visual field. We started by monocularly presenting local motion within circular patches in front of a static visual background across the zebrafish’s monocular visual field to record spatially resolved calcium responses in the contralateral optic tectum (Figure 2A). Stimuli were designed to mimic local motion in visual subfields that fish would usually perceive during backwards egomotion (Figure 2A, red arrows). We used zebrafish expressing GCaMP6f pan-neuronally and recorded from a total of 17,844 ROIs in the optic tectum. To estimate the neuronal receptive fields (RFs), we first calculated the weighted activity maps based on stimulus location, stimulus size and the cell’s calcium trace (Figure 2C top), and then isolated the spatial receptive field (RF) using agglomerative clustering (Figure 2C bottom). We indicate the RF size in steradian: one steradian (sr) equals a solid angle of 65° (or 3,282 square degrees). The RF sizes obtained by this were organized in two large clusters (S-Figure 2B), one (n = 11,240 ROIs) with sizes close to the maximum extent of the stimulated area (6.18 +/- 0.47 steradian) - with generally low inter-trial reproducibility (0.06 +/- 0.11) - and another (n = 6,604) with smaller sizes (0.48 +/- 0.37 sr) and larger reproducibility of responses (0.47 +/- 0.27). For our further analysis, we removed RFs that were larger than 2.0 sr and ROIs whose responses varied substantially between repeated presentations of the same stimulus (reproducibility index < 0.3). RF sizes for cells passing these criteria (n = 4,745) appeared to follow a Weibull distribution (Kolmogorov–Smirnov test, D = 0.0172, p = 0.1198, Figure 2D). The RF center locations in retinocentric visual space were distributed across the whole monocular visual field of view of the eye contralateral to the recorded tectal hemisphere. Notably, RF densities (Figure 2E) for the forward moving stimuli appeared considerably higher in the temporal visual field (i.e. nasal locations on the retina), which corresponds to the observation that the visual stimuli moving towards the animal evoke larger responses than outward moving stimuli (Naumann et al., 2016), but is in apparent contrast with the higher photoreceptor density covering the nasal visual field (Zimmermann et al., 2018).

**Figure 2.**
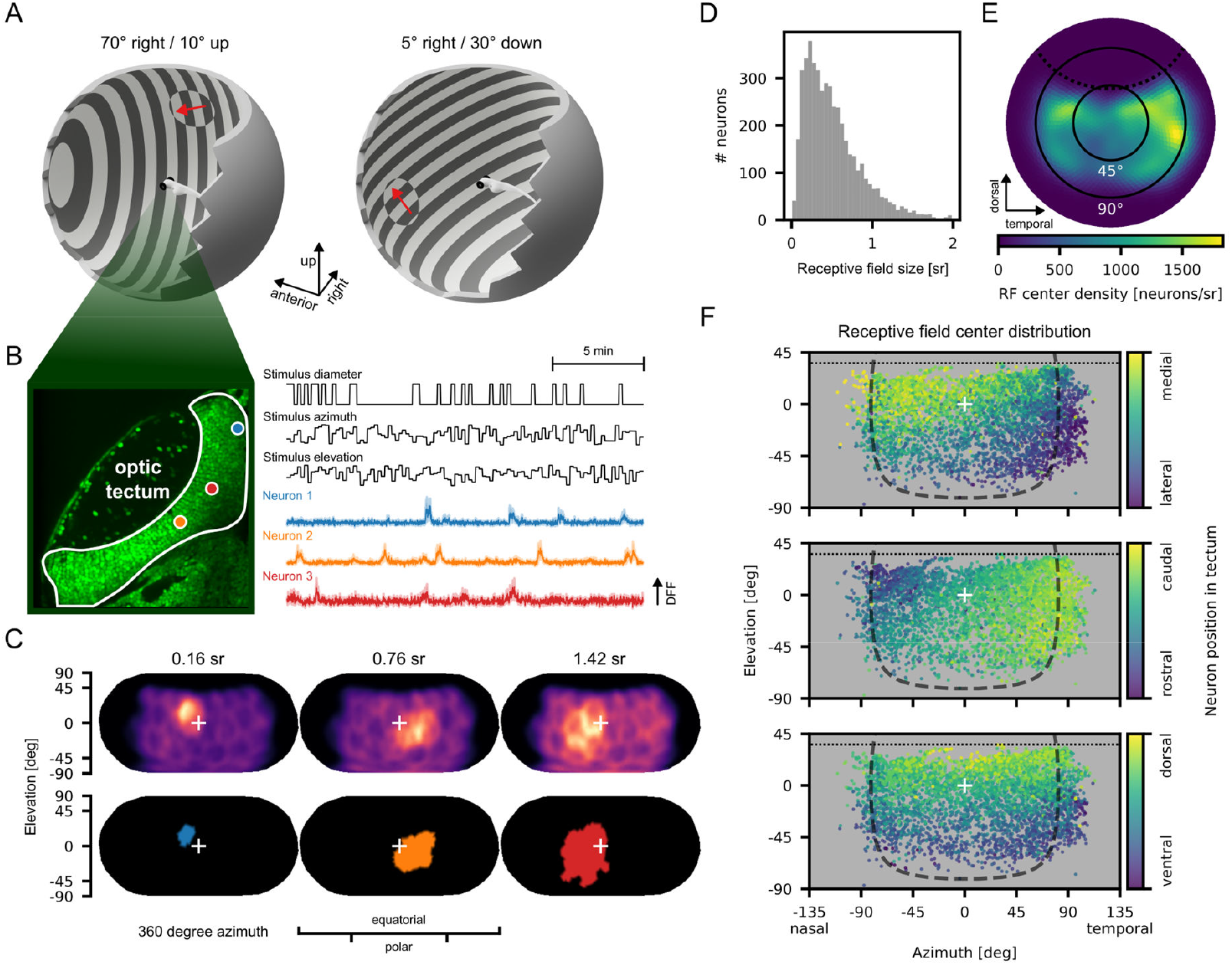
Spatial receptive (RF) field mapping of motion-sensitive cells in the optic tectum (n=7 fish, 5-7 dpf). **A**: Stimulus example. Fish were monocularly presented with local motion gratings at different locations in visual space in front of a static background grating. The motion direction was always in the direction of a backward body translation, and gratings were oriented perpendicular to the motion direction. **B**: 2-photon recording of GCaMP6f in a layer of optic tectum with three example cells highlighted (blue, orange, red) and their corresponding mean activity across repetitions during visual stimulation plotted over time. **C**: Retinocentric weighted activity maps across visual space (top) for example units in B and corresponding segmented receptive field (RF) maps (bottom). White cross marks the estimated center of the retina. Data is shown using Eckert IV equal-area projection **D**: Distribution of RF sizes for all neurons included in the analysis. **E**: Retinocentric visual space projection of the density (neurons/steradian) of RF centers across the monocular visual field. Radial directions are dorsal (top), temporal (right), ventral (bottom) and nasal (left). Distances to the center (white cross) of the projection are in degrees of visual angle. Dotted black line marks the border above which the animal could not be stimulated because of the microscope objective obstructing the view. **F**: Distribution of RF barycenters across right eye retinocentric visual space with relative anatomical position color coded: medio-lateral (top), rostro-caudal (middle) and ventro-dorsal (bottom). Dashed black line shows estimate of monocular visual field according to (Easter, Jr. & Nicola, 1996). Dotted black line as in panel E.

The relatively dense accumulation of RF centers in the temporal visual field of fish initially led us to suspect that possible optical artifacts from the direction of the transparent mounting rod (Figure 1D c) might have skewed the results of our analysis. To mitigate this potential issue, in 3 of our 7 fish, we conducted the experiments while they were embedded at a 90° horizontal angle, thus completely avoiding stimulation around the mounting base. The data from these measurements were virtually identical to those in the standard embedding, thus showing that the results are not stimulation artefacts, and are already included in Figure 2D to F.

To our surprise, the distribution of tectal RF centers across the fish’s retinocentric visual field shows that the overall extent of the visual field exceeds the previously suggested monocular visual field extent of 163° in larval zebrafish at 96 hpf (Easter, Jr. & Nicola, 1996, dashed black line) – particularly in the temporal visual field. RF centers could be found at elevations down to roughly -70° in retinal elevation. In combination with the RF sizes (Figure 2D), this finding suggests that the RF area can also touch the nadir (the region directly below the fish), although the recorded fish sampled this region less densely than other regions of visual space. Registration of anatomical cell centers within the tectum showed a clear topographic gradient (Figure 2F) in line with previously reported findings in tectal recordings (Holt, 1983; Wang et al., 2020) - cells with RFs in the upper nasal visual field were found in more medio-rostral parts of the dorsal tectum, while those with RFs in the lower temporal field, were more prevalent in latero-caudal parts of the ventral tectum. Taken together, our tectal recordings suggest a visual field of +/-90° in retinal azimuth and down to -70° in retinal elevation. These measurements are relative to the eye’s center axis, which at rest is pointing slightly upward [+4° according to Dehmelt et al. (2021)] and is slightly converged horizontally (between 7 and 17° toward rostral in all measured fish) in body space coordinates. These results indicate that, depending on the deflection of both eyes, larval zebrafish are in principle able to observe the whole horizontal extent of their environment.

### Global-flow responsive cells sample visual field down to nadir and show bias for lower visual field in pretectum

Visuomotor stabilization responses such as the optomotor response are mediated by global-flow responsive neurons. These neurons are only sub-optimally stimulated by displays positioned on the side (Zhang et al., 2022) or below the animal (Kist & Portugues, 2019), because of their often large RFs. We therefore used a contiguous motion noise (CMN) stimulus (Figure 3A top left, Zhang & Arrenberg, 2019) in our MARINER arena to map the entire extent of RFs, that is the local motion preferences of each cell in different parts of its RF across the complete visual field. The CMN stimulus provided random, but locally coherent and globally contiguous visual motion stimuli that are amenable to slow 2-photon calcium imaging when using reverse correlation techniques for RF estimation (Figure 3A).

**Figure 3.**
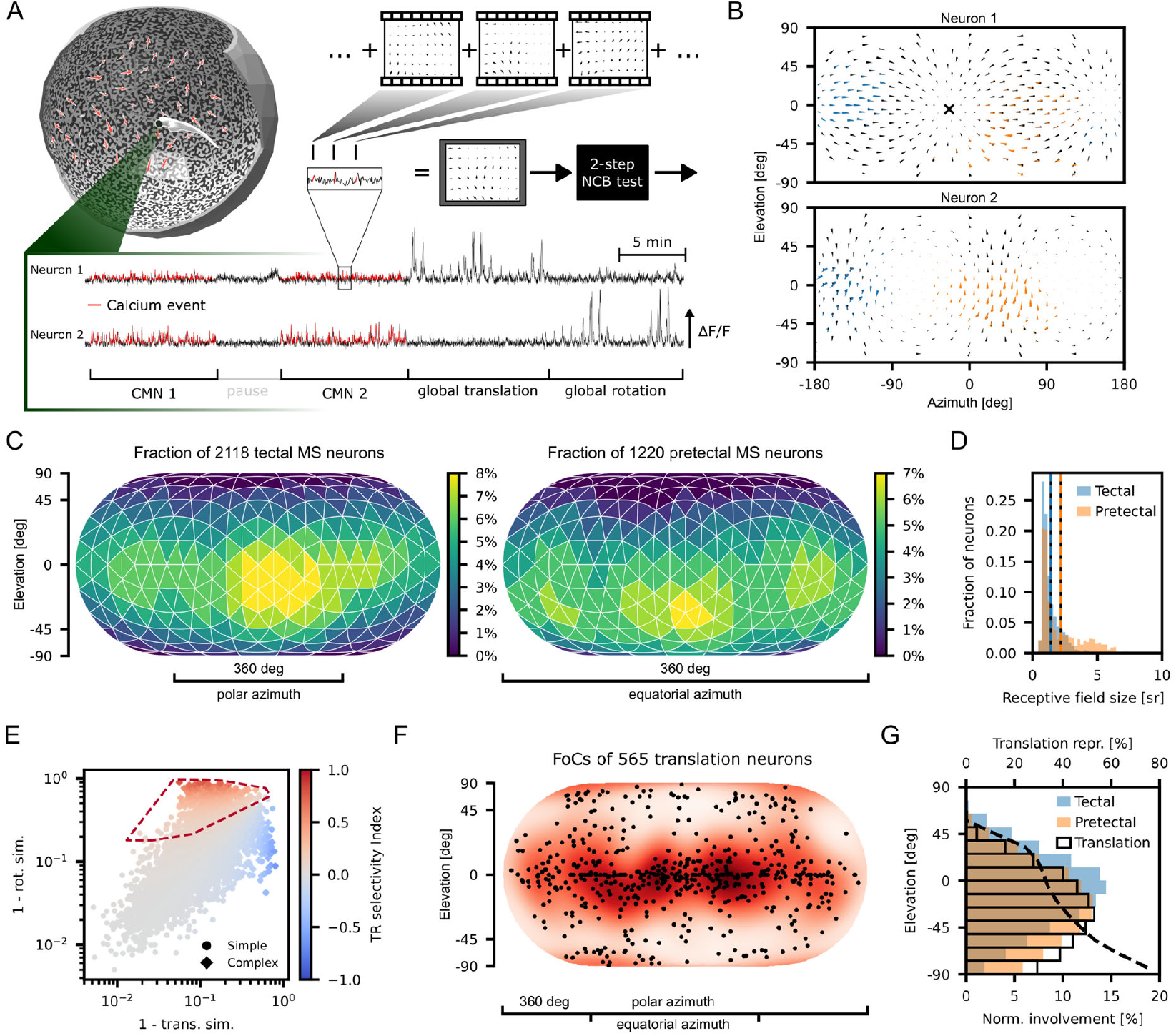
Motion receptive field (RF) mapping of cells in the optic tectum and pretectum (n=4 fish, 5-7 dpf). **A**: Fish were presented with a contiguous motion noise stimulus (CMN, top left, S-Video 2) during 2-photon calcium imaging of the mid-and forebrain. Motion RFs were calculated using a reverse correlation estimate during calcium events and validated using a cluster-based bootstrap test described previously (2-step NCB test, Zhang & Arrenberg, 2019). **B**: Example quiver plots of the RFs of a translation (top) and rotation (bottom) selective cell in the pretectum. Black arrows show the best ego-motion template match for a given cell. Colors show spatially separate subfields of its RF and their preferred local motion direction. Black cross for neuron 1 marks the RF’s focus of contraction (FoC) of the translation selective example neuron 1. **C**: Heatmap of the fraction of cells for which the respective locations in the fish’s visual space were part of its RF for tectal (left) and pretectal (right) cells. Data is shown using Eckert IV equal-area projection in body-centered coordinates of visual space **D**: Distribution of receptive field sizes separated in color by anatomic cell location, with means marked by vertical lines for tectum (blue) and pretectum (orange). **E**: Best rotation and translation similarities plotted against each other with TR selectivity index color coded. Dashed line shows convex hull of all translation-selective cells (TR index > 0.1) selected for panel G. Symbols show simple and complex cells (with one or more than one RF subfield, respectively) **F**: Map of the foci of contraction (FoC) for all translation selective cells (black dots). Red heatmap shows a relative density estimate of the FoC distribution (neurons/steradian). **G**: Histogram of relative brain area involvement shown in C based on elevation. Bars are normalized to account for elevation-dependent differences in area on sphere. Black dashed line shows the relative translation cell representation given the overall tectal and pretectal involvement at each elevation: i.e. the ratio of cell counts for translation RFs covering this elevation divided by the number of all tectal and pretectal cells covering this elevation.

Out of 36,359 ROIs in 6 fish (5-7 dpf), we identified 5,154 motion-sensitive neurons, primarily in the mid- and forebrain. Of these, 2,118 were in the tectum and 1,220 in the pretectum. Analysis of the contributions of different locations in visual space towards the neurons’ total RF showed that tectal cells are mainly represented in the equatorial regions, whereas pretectal cells preferred somewhat lower elevations (Figure 3C and G). Notably, the identified pretectal receptive fields covered the region around the nadir almost as well as other regions of visual space, suggesting that the animals also use stimuli directly below them to inform stabilization behaviors. In the optic tectum, on the other hand, the region of the nadir is sampled more scarcely.

RFs in pretectal cells were on average larger (2.18 +/- 1.65 steradian) than tectal cells (1.41 +/- 0.80 steradian), and the pretectal RF size distribution was right-skewed (Q3 3.10 steradian in pretectum vs. Q3 1.69 steradian in tectum, Figure 3D). RFs were not always contiguous but consisted of one to three spatially separate subfields (modes). To differentiate between cells that were motion-sensitive and those who preferred specific types of egomotion, we calculated a translation/rotation selectivity index (TR index) for each neuron, based on their best cosine similarity to egomotion templates. Cells with one subfield (simple cells) generally had TR indices closer to zero (S-Figure 3A), while those with two or three subfields (complex cells) were more likely to deviate towards more positive or negative values (indicating translation and rotation selectivity, respectively). In line with previous results (Zhang et al., 2022), and indicative of their smaller RFs, simple cells also had higher overall cosine similarities to their preferred egomotion template than complex cells (S-Figure 3B). Both translation- and rotation-selective cells were present in our dataset, and a much higher proportion of rotation cells were found than in the previous study. To further compare our results to previous findings, we selected cells with more translation-like characteristics by filtering for any cells with a TR index larger than 0.1 Figure 3E, circumscribed area with dashed line). The distributions of their foci of contraction (FoC) were qualitatively identical to previously described results (Zhang et al., 2022), with a major concentration around the equatorial region, lesser concentrations at the poles and with few neurons encoding oblique translation axes (Figure 3 F). Interestingly, we found that translation cells were strongly over-represented at lower visual elevations (Figure 3G, black edged bars and dashed line), which is in line with predictions about optimal decoding of translational optic-flow made based on natural scene statistics (Alexander et al., 2022).

### Motion vision is red-green colorblind in tectal cells and during optokinetic response

To demonstrate MARINER’s capability for concurrent multi-spectral stimulation, we investigated color processing in the midbrain. Color-opponent processing channels have been shown to exist in the retina (Zhou et al., 2020), while the optomotor response (a gaze stabilization behavior) has been reported to be color-blind in zebrafish (Orger & Baier, 2005). This result suggests that motion-sensitive neurons in the zebrafish brain are colorblind, although direct neurophysiological evidence is missing. We recorded calcium activity in the optic tectum of three fish while presenting whole field moving color contrast gratings in which the phase and antiphase were varied in red (625 nm) and “green” (475 nm) contrast, respectively (Figure 4A top and 4D). If information from the L cone and the M cone were separately maintained in the downstream visual circuits, we would expect red/green color-opponent circuits and strong responses to a moving pattern of alternating red and green stripes. However, Orger and Baier (2005) suggest that at least the circuits underlying the optomotor response merge the red and green channels in the motion vision pathways, such that responses to alternating red and green stripes are small or absent, i.e. red-green colorblind for optomotor responses. After correlation analysis with red, green and achromatic regressors (S-Figure 4A) for clockwise (CW) and counterclockwise (CCW) direction selectivity and non-directional motion sensitivity, we identified 45 (CW), 55 (CCW) and 73 (MS) cells for each category (Figure 4B). Responses of direction-selective (DS) cells were attenuated for red-green antagonistic stimuli (Figure 4E) when compared to monochromatic stimuli evoked by only one of the LEDs (Figure 4E left column and bottom row). Responses were especially low when the red LED was about 1.8x brighter than our green LED. Additionally, in 2 animals, we tracked eye positions during stimulation to assess their optokinetic response (OKR) performance (Figure 4C). The color contrast tuning of OKR slow phase was qualitatively similar to the tuning of DS cells (Figure 4F). Both OKR behavior and tectal DS cells responded strongly to high monochromatic contrasts (upper left and lower right corners in Figure 4D-F), with fish gradually decreasing their stimulus-driven responses as luminance contrasts approached zero (approx. diagonals in Figure 4D-F). Together, our results show that tectal DS cells and the OKR appear to be red-green colorblind. Interestingly, we observed some notable differences between the responses of MS and DS cells. MS cells appeared to be more strongly driven by red light stimulation, whereas most DS cells were driven by both red and green light (Figure 4G-H). DS cells also seemed to show a larger range of graded responses to (achromatic) luminance modulations (Figure 4I), while MS cells responded primarily to the higher contrast levels. Many of the non-DS cells that were excluded from analysis here showed higher diversity regarding their chromatic response types (S-Figure 4A), suggesting that chromaticity may play a more important role in other types of visual neurons in the optic tectum (Fornetto et al., 2020; Guggiana Nilo et al., 2021).

**Figure 4.**
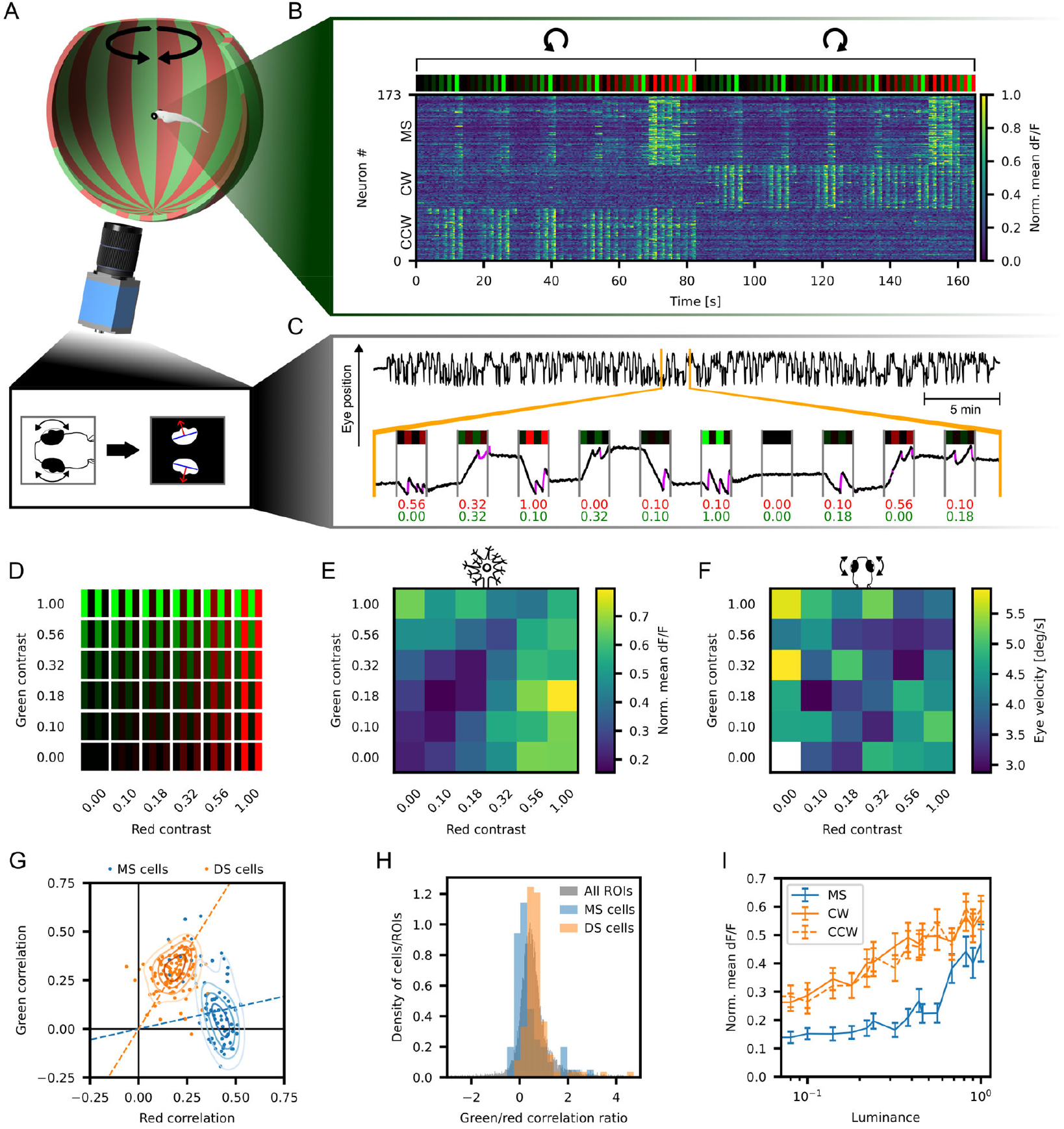
Behavioral and tectal responses to color-opponent global rotations. **A**: Stimulus example. Fish (n=3, 5-6 dpf) were visually stimulated with color opponent gratings, while recording neuronal activity and eye positions. **B**: Raster plot of repetition-averaged, normalized dF/F for all global motion and direction-selective cells in the optic tectum (out of total of n= 17,364) in response to clockwise and counterclockwise global moving gratings. Shown are motion-sensitive (MS, n=62), clockwise (CW, n=32), and counterclockwise (CCW, n=43) direction-selective cells. **C**: Example eye position recording (top). Inset (bottom) shows zoomed in segment with moving stimulus phases marked. Number annotations display uncalibrated, relative red and green intensity levels during stimulation phases. Light purple segments on eye trace mark saccade instances, which were discarded for further analysis. **D**: Illustration of the color contrast combinations that were shown in phase and antiphase of the binary gratings. **E**: Summary data for normalized mean dF/F of 100 direction-selective cells (CW and CCW) in 3 fish as a function of color contrast combinations shown in D. **F**: Summary data for mean eye velocities of left and right eyes in 14 recordings from 2 fish. **G**: Correlation coefficients to the red and green plotted with cell identity (direction-selective, DS, or motion-sensitive, MS) color coded. Colored dashed lines are lines going through origin and cell type’s median correlation coordinates. For each DS cell the corresponding directional type’s regressor correlation is plotted (i.e. CW regressor correlation for CW cell and the same for CCW cells). **H**: Density distribution of green/red correlation ratios for MS and DS cells and with all segmented ROIs as reference. For all ROIs (gray) the highest regressor correlations out of all (MS, CW, CCW) were used. **I**: Normalized mean dF/F for each cell type as function of achromatic contrast (luminance). Error bars show SEM.

## Discussion

We present a visual stimulation arena that is suitable for small aquatic species and enables unprecedented control over spatiotemporal and chromatic stimulus features at the same time and across most of the animal’s visual field. MARINER avoids common problems of underwater visual stimulation that can be caused by refraction and reflection (Dunn & Fitzgerald, 2020; Wang et al., 2021). Previously published stimulation solutions often succeeded in improving individual stimulus features, such as increasing visual field extent through spherical geometry (Dehmelt et al., 2021), or reducing optical artifacts through stimulus back projections (Huang et al., 2020; Stowers et al., 2017), while neglecting features of lower importance to the particular experiment, like chromaticity or visual field coverage, respectively. MARINER is designed for concurrent 2-photon calcium imaging and behavioral recordings during visual stimulation. Since all visual stimuli are rendered in real-time through a Python/OpenGL based software (Hladnik, 2025), it is also possible to use the arena for closed-loop experimental designs (see for example saccade triggered stimulation in Soto et al., 2025).

Our spatial RF mapping results show that the total monocular visual field of larval zebrafish encompasses more than the previously proposed 163° azimuth (Easter, Jr. & Nicola, 1996), pointing to the special design challenges and subsequent adaptations of biological lenses for underwater vision, which typically use a gradient refractive index composition (Jagger, 1992; Young et al., 2018). Our motion RF experiments confirmed and extended previous observations (Zhang et al., 2022) about the directional characteristics of translation-selective cells that are primarily found in the pretectum (Figure 3F). MARINER’s ability to stimulate down beyond the previous limit of -40° elevation enabled us to demonstrate a distinct representational bias (Figure 3G) among these cells for low elevations close to nadir, that had previously been proposed on the basis of natural scene motion statistics (Alexander et al., 2022) and behavioral experiments (Kist & Portugues, 2019; Wang et al., 2020).

The chromatic resolution of MARINER allowed us to collect neurophysiological and behavioral evidence on colorblindness of one particular behavior, the optokinetic response. Other behaviors and circuit pathways are expected to have different chromatic preferences [e.g. UV light for prey capture, Khan et al. (2023)] and may exploit the information contained in color-opponent retinal ganglion cells (Zhou et al., 2020). Since the last common ancestor of mammals and teleosts was probably tetrachromatic (Baden & Osorio, 2019) and the loss of tetrachromatic vision in mammals is likely attributable to a “nocturnal bottleneck” in mammalian evolution (Gerkema et al., 2013), the study of aquatic vision in teleosts promises insights regarding circuit design and adaptations of color vision in vertebrates. Further work is needed to reveal the relevance of color opponency (Zhou et al., 2020) and the contributions of each cone channel to behaviors like prey capture (Yoshimatsu et al., 2020) or escape and the various functionally identified cell types in the visual brain (Heap et al., 2018; Wang et al., 2020).

Our hope is that MARINER will help to unify different parts of aquatic vision neuroscience research, by providing a platform that enables spatially well resolved, polychromatic visual stimulation in combination with neurophysiological and behavioral measurements.

## Methods

### Animal handling

All animal work conformed to German laws for animal research and was in accordance with the local governmental guidelines (Regierungspräsidium Tübingen). Larval zebrafish (*Danio rerio*) transgenic for *Tg(HuC:H2B-GCaMP6f)jf7Tg* and homozygous for the nacre mutation were used for all 2-photon imaging and behavioral experiments at 5 to 7 days-post-fertilization.

### In-vivo 2-photon calcium imaging

For imaging experiments, fish were embedded on a Petri dish in 1.7% low-melting agarose and transferred onto a stage made of 2.5% agarose - bonded to the plastic arrowhead (Figure 1Db, S-Figure 1A). The pipette tip was then mounted in the center of the spherical enclosure of the stimulation arena by sliding it into a hollow glass rod (Figure 1Dc) protruding towards the center of the enclosure. The container was then filled to the top with E3 medium. For experiments where tracking of eye movements was required, the eyes were cut free of agarose using a custom platinum wire tool (Arrenberg, 2016).

Functional fluorescence imaging of the genetically encoded calcium indicator GCaMP6f was done using a galvo-galvo movable-objective 2-photon microscope from Sutter Instruments (Novato, California, USA) with a Coherent Vision-S Ti-Sa laser set to 920nm and a 20x/1.0 Zeiss water-submersion objective (APOPLAN). Image time series were acquired one dorso-ventral layer at a time at a framerate of 2.18 Hz and with a resolution of 512 × 512 pixels using Sutter Instruments’ MScan software.

### Visual stimulation and behavioral imaging

Visual stimulation was done using a single light-guide-coupled DMD video projector (Lightcrafter E4500 MKII, model: DPM-E4500MKIIFC-OX, EKB technologies) to cover roughly 360° azimuth and 135° elevation on a 3D printed spherical enclosure (“fishbowl”, material Accura ClearVue or Formlabs BioMed Clear) that served as both water container and back-projection screen. The outside of the fishbowl was coated in white acrylic paint (Tamiya Color XF-2) to increase the diffusive properties of the surface, and the coating was protected from mechanical abrasion by applying a clear acryl-based varnish. The projection was achieved by splitting the first focused image of the video projector on a 4-sided pyramidal mirror (side slopes 22°), then re-focusing and magnifying each sub image through a 75 mm focal length biconvex achromatic doublet lens (mounted at 44°), and folding the light from each of the 4 sub images in the direction of the centrally mounted fishbowl (via a 78° mirror) with the fish mounted in agarose in its center. The video projector was powered by a multispectral high power LED light source (CHROLIS, 365/420/470/565/590/625 nm, Thorlabs).

The generation of images for visual stimulation and the acquisition of behavioral tracking data were performed using the custom written Python/OpenGL software VxPy (https://github.com/thladnik/vxpy). Tracking of eye positions was made possible by infrared back-illumination through the microscope’s objective and a transparent, circular microscopy coverslip glued to the bottom of the spherical enclosure. Through this coverslip images from the zebrafish larva could be captured by a 45° folding mirror mounted directly below the enclosure and were recorded by a camera (Basler ace 2 camera a2A1920-160umBAS) at up to 2.3MP and 160 Hz (S-Figure 1C).

During visual stimulation with concurrent 2-photon calcium imaging, the light emission of the LED light source was blanked intermittently in synchrony with line scans. Light for visual stimulation was only emitted for the duration of the Y-mirror flyback to avoid contamination of the imaging. At the same time, the photomultiplier tubes were turned off by a second, gating signal, to avoid overexposure and damage. We have outlined a simply blanking circuit using a Teensy 4.1 microcontroller for Galvo-Galvo and Resonant-Galvo scanners in S-Figure 1D.

A full list of parts as well as the 3D design files for customized parts and necessary software for LED blanking are freely available at the MARINER repository (https://github.com/thladnik/mariner-experimental-setup).

### Coordinate-space conventions

Unless stated otherwise, stimulus coordinates, when expressed in spherical coordinates, use spherical polar coordinates on a unit sphere. The coordinate system is fish-centered, with 0° elevation and 0° azimuth corresponding to the rostrum (caudo-rostral axis). Elevations are normalized to the range -90 to +90° with positive angles moving dorsally and azimuths are normalized to the range -180° to +180° with positive angles moving rightward. When expressed in 3D cartesian coordinates, we use a right-handed coordinate system, where the positive x, y and z directions correspond to the fish’s caudo-rostral, leftward, and ventro-dorsal directions, respectively. Note that this notation leads to a sign switch during transformation from azimuth to y-axis.

For clarity, elevation Θ and azimuth ϕ may be calculated from cartesian coordinates (x, y, z) as

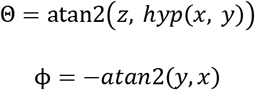

while the reverse is possible with

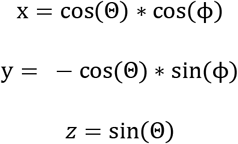

In cases where area-preserving projections were necessary for illustrative purposes, we used Eckert-IV equal-area projections (Snyder & Voxland, 1989). Cartesian 2D coordinates (x, y) were calculated based on the spherical coordinates as

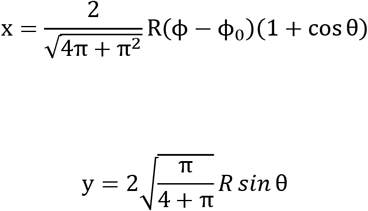

Where ϕ_0_ is 0 and θ is the auxiliary angle, which is the iteratively determined solution to

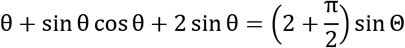

using the Newton-Raphson method.

### Calcium imaging data preprocessing

MScan MDF imaging data files were converted to TIFF file format for further analysis. The imaging time series data was registered and segmented and the raw fluorescence calcium signal for each segmented region of interest (ROI) was extracted based on anatomy using Python 3.10 and the suite2p package (Pachitariu et al., 2016). Based on the raw fluorescent signal sum of each ROI, the instantaneous change in activity dF/F for each imaging frame was calculated as

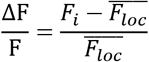

where *F*_*i*_ is the sum fluorescent signal of the ROI in the *i*-th imaging frame and *F*_*loc*_is the local arithmetic mean of all values in the 10th percentile out of all the fluorescent signals within a 120s time window centered on the *i*-th imaging frame. This approach ensured a dF/F signal with little to no change in the baseline dF/F (i. e. in absence of elevated neuronal activity) throughout the whole recording, despite continuous decrease of the mean fluorescence due to bleaching.

### Anatomical registration to reference atlas

ROI centers obtained from suite2p were mapped to the MapZeBrain atlas (Kunst et al., 2019) via a 3-step transform using the Python bindings to the Advanced Normalization Toolbox ANTs (Tustison et al., 2021). First, the mean image of a recorded layer was registered to a z-stack of the recorded brain area (local stack) using an affine transformation. Second, the local stack was registered (SyNRA) to a z-stack of the whole brain for the transgenic *Tg(HuC:H2B-GCaMP6f)jf7Tg* line (standard stack). Lastly, the standard stack was mapped (SyNRA) to the *jf5Tg[Tg(elavl3:Hsa*.*H2B-GCaMP6s), HuC:H2bGCaMP6s, elavl3:H2bGCaMP6s]* marker line of the MapZeBrain atlas. In cases where cells were characterized based on their anatomical localization in particular brain areas, the registered 3D cell positions were compared to the atlas’ region data (version 2.0, MECE 2024).

### Estimation of spatial extent of motion receptive fields in optic tectum

#### Visual stimulation

Larval zebrafish were mounted for 2-photon imaging as described above. Fish were (mostly) monocularly stimulated with red light (625nm, typical power 490mW) while recording from the contralateral side of the optic tectum. They were presented with a stimulus protocol consisting of a static binary grating background pattern with a spatial frequency of 0.05 cyc/deg and localized circular patches at different locations in the fish’s visual field wherein the binary grating was moving at 30 deg/s. The orientation of the grating and direction of the motion were set to locally mimic forward optic flow, meaning the direction of motion was towards a focus of contraction directly rostral of the fish (in positive x-direction), with the grating oriented perpendicular to the direction of motion (see Fig. 2A). Stimulus patch centers were evenly distributed across the visual field, at locations that were determined by a repulsive sphere model, which simulates repulsive interaction of a fixed number of such centers based on the geodesic distances between centers in order to achieve center distributions with uniform density across the sphere (Dehmelt et al., 2021). In case of stimulation of the right eye, for example, stimulus centers were constrained to cover the whole right side of the visual field, including locations up to -40° azimuth in front of the fish and -15° azimuth behind the fish, for a total coverage of 235° azimuth. Elevation-wise, stimulus centers covered the maximum range - a total of 135° from -90°deg to +45°.

Two different patch sizes were used for visual stimulation. Small patches with 25 deg diameter were presented at 78 different locations across the visual field and large patches with 60 deg diameter at 24 locations (S-Figure 2A). For each stimulus presentation, fish were shown the static background grating phase for 4s (motion velocity 0 deg/s), followed by 4s phase where the grating within the stimulus patch was moving at 30 deg/s. The stimulus presentation order was randomized, and each stimulus location and patch size combination was presented three times, resulting in a total of (78 + 24) * 2 * 3 = 612 stimulus phases per recording.

#### Analysis

For each ROI we calculated the mean dF/F during any given stimulus phase. This resulted in a 3 × 204 array wherein each element corresponded to the activity for a given stimulus location, patch size and stimulus velocity during one of 3 stimulus repeats (or motion stimulus pauses). From this we calculated the mean signal autocorrelation between repeats, which indicated the reproducibility of the ROI’s response during repeats (“repeatability”). Based on the distribution of autocorrelation coefficients, we set a lower threshold of 0.4 to filter for ROIs that responded consistently to repeat presentations of the same visual stimulus sequence. For each ROI that passed the repeatability threshold, we calculated a weighted mean stimulus map. We did this by creating a sphere of 2000 evenly distributed points (see above) and then calculating a binary mask, wherein each point was assigned a value of either 1 or 0, based on whether the stimulus motion was presented at this point or not. This resulted in a 2000 × 306 array of 0s and 1s, where each point within the stimulation patch was assigned a 1. We then calculated the weighted mean stimulus map by weighing each point’s contribution based on the ROI’s mean dF/F during any of the 306 moving stimulation phases, resulting in a 2000 entries long vector of relative dF/F activity at the 2000 points on the sphere. The resulting weighted map was then smoothed, by convolving it with a 3-dimensional von-Mises-Fisher function *C*3 with a concentration parameter κ of 150:

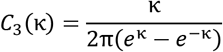

The relative activity values on the smoothed map were then classified using an agglomerative clustering algorithm with 3 components and a cosine distance constraint, where one component represented the response to un-stimulated parts of the visual field, a second one the background activity outside the receptive field (RF) and a third the activity inside the RF. The component with the highest relative mean activity was then selected as the RF. From the total number of points that were classified as part of the RF, we calculated the RF size in steradian by multiplying the number of points select as part of a RF with the solid angle subtended by a single point (0.00628 sr/point). Based on the corresponding point positions on the sphere that were classified as part of the RF, we calculated the 3D barycenter of the RF.

### Estimation of local motion receptive fields in optic tectum and pretectum using the CMN stimulus

#### Visual stimulation

Larval zebrafish were mounted for 2-photon imaging as described above. For stimulation, 625nm (typical power 490mW) and 475nm (typical power 630mW) light was used. Larvae were presented with a full field Contiguous Motion Noise (CMN) stimulus as described previously (Zhang et al., 2022; Zhang & Arrenberg, 2019) with a few notable alterations to account for the spherical geometry the stimulation setup outlined below.

To keep the individual stimulation patches that displayed the moving binary noise patterns approximately uniform in size across the whole sphere, they were constructed to be triangular – as opposed to the original quadratic shape – based on a subdivided icosahedral body with 320 faces. The binary noise texture itself was generated based on the cartesian position unit vector coordinates and a 3D simplex noise algorithm described by McEwan et al. (2012), instead of the original mapped 2D uniform distribution. The CMN stimulation parameters were chosen to match the velocity statistics of the original implementation in Zhang & Arrenberg (2019). Two segments of the CMN stimulus, 10min each, were presented separated by a 5min pause without stimulation. The CMN stimulation was followed by a translation and rotation characterization protocol, consisting of global gratings with a period of 20° moving at a speed of 30°/s. Gratings were moving along or around one of 32 axes of different azimuths and elevations, either in positive or negative direction (translation) or in clockwise or counterclockwise direction (rotation). Each 4s moving stimulus phase was preceded by a 4s static phase with gratings in the same direction as the moving phase and each motion type/direction combination was presented 2 times.

#### Analysis

Analysis of data from CMN experiments was performed in the same way as outlined in Zhang and Arrenberg (2019) with the following alterations. The 2-step non-parametric cluster-based bootstrap (2-step NCB) analysis was performed in Python instead of Matlab. The Calcium event instances were calculated based on a smoothened derivative of the cell’s dF/F and a fractional threshold on the dF/F’s magnitude. Also, because of the triangular geometry of stimulus patches, in the 2^nd^ step, for the tracing of patch clusters with significant responses in the 1^st^ step, only the three nearest patches were considered, instead of the original four.

### Estimation of differential chromatic contributions to global motion processing in tectal cells

#### Visual stimulation

Fish were stimulated with a global moving grating stimulus with a spatial period of 30 degrees that was either static or moving at 30 degrees/s in clockwise or counterclockwise direction. The two phases of the optical grating were displayed in different colors. One phase of the grating was illuminated using the 625nm LED (“red”, typical power 490mW) and the other using the 475nm LED (“green”, typical power 630mW) – both a varying relative power levels of 0.00, 0.10, 0.18, 0.32, 0.56 and 1.00 – making for a total of 36 individual stimulation combinations per trial repeat and stimulus direction. Each 4 second moving stimulation phase was preceded by a 4 second static phase with the same relative red-green contrast ratios and each individual combination was shown 3 times – resulting in a total of 432 stimulation phases. Combinations were displayed in the same pseudorandomized order for each of the 3 repeats.

#### Analysis

##### Tectal activity

ROIs were filtered based on their inter-repeat reproducibility (“repeatability”) as described above. For further analysis only cells with a repeatability larger than 0.3 were considered. Next, the correlations of cells’ dF/F in response to different motion regressors were calculated. We constructed one regressor each for red, green and achromatic response types and for each possible motion response type: motion-sensitive (MS), clockwise direction-selective (CW) and counterclockwise direction-selective (CCW), leaving us with a total of 9 different regressors and corresponding correlations for each cell (compare S-Figure 4A). From these we selected the exclusively MS, CW and CCW selective cells by applying a universal regressor correlation threshold of 0.35. Meaning that for example, for cells to be considered CW selective, they must have a CW correlation > 0.35, an MS correlation < 0.35 and a CCW correlation < 0.35 for either red, green, or achromatic regressors. For each of the 36 contrast combinations we then calculated the mean dF/F based on the between-repeat mean dF/F values.

##### Eye movements

We calculated the instantaneous eye velocity from the eye position data. We used this velocity data to identify saccades by using a 3-component gaussian mixture model to identify the highest velocity component and exclude all its data points from further analysis. Based on the remaining velocity data we calculated first the per-phase mean eye velocity values and from there the per-contrast condition mean eye velocity.

### Irradiance measurements and contrast calculation

To estimate the maximum possible local contrast for visual stimulation we measured the radiant flux at different elevations for each stimulus color channel. Measurements were performed using a Newport 1918-R optical power meter with a 918D-ST-UV photodiode probe and the attenuator disabled. The sensor was positioned directly on the inside wall of the fishbowl, while projecting a checkerboard pattern with a 90° period in azimuth and elevation. For each brightness level two measurements were taken (S-Table 2): one on a bright patch of the checker pattern (E_max_) and one on the dark patch (E_min_). Power readings in μW were converted to irradiance based on the surface area (A) of the sensor with

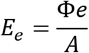

where *E*_*e*_ is the irradiance and Φ*e* the measured radiant flux. Michelson contrasts *C* (S-Table 3) for each color channel separately were calculated as

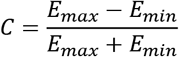

### Spatial resolution measurements using Siemens stars

For the calculation of the spatial resolution limits (S-Figure 1E), Siemens star patterns with a diameter of 20° and a spatial period of 45° were projected in a northern, southern, western and equatorial location (top left) using a monochrome Basler a2A3840-45umPROcamera with a 4k resolution. The cropped-out star pattern was then rotated in-silico at 90°/s using Python and OpenCV (bottom left) and a temporal resolution power spectral density was calculated for each camera pixel (bottom right). The mean pixel power spectral density for each Siemens start location was then used to infer a spatial resolution limit.

### Limitations of MARINER

Despite its advantages, some persistent technical challenges remain. Most notably, as with any other approach that tries to combine visual stimulation with optical imaging techniques, light pollution of the imaging poses a significant problem. MARINER solves this by intermittent blanking of the visual stimulation during signal acquisition – either synchronized to the mirror flyback periods or to the imaging frame rate in Galvo-Galvo and Resonant-Galvo scan mode, respectively (see S-Figure S1D). However, this can often lead to temporal aliasing – particularly at low contrast levels when the overall illumination times are shorter due to temporal light intensity modulation of the video projector output. Some ways to ameliorate this problem would be to use multi-DMD systems to reduce the prevalence of short stimulation time windows for individual color channels and to use optical filtering of stimulation light in combination with programmable denoising techniques based on prior knowledge of temporal stimulus luminance data to improve signal quality. Another technical challenge is the achievable contrast level in any full-surround arena. Individual photons can briefly get trapped in full-surround arenas, which essentially operate like Ulbricht spheres (Banda, 1999) and the escape of the photons (which would increase contrast) depends on the reflection coefficient of the inner surface of the sphere. This relation suggests that higher contrasts should be achievable with gray (not white) screens, at the expense of overall brightness.

## Supporting information

MARINER light path schematic drawing

Bill of materials

Stimulus arena animation

Visual stimulus examples

Illustration of Figure 2 receptive field mapping process for population activity

## Acknowledgements

We would like to thank Hansjürgen Dahmen for his help in conceptualizing the optical design of the visual stimulation arena, and Carina Thomas and Giulia Soto for all the fish. We would also like to acknowledge the PennState BioAtlas (https://bio-atlas.psu.edu/, NIH grant 5R24 RR01744, Jake Gittlen Cancer Research Foundation, and PA Tobacco Settlement Fund) from which the anatomical zebrafish slices for S-Figure 2 were taken. This work was funded by the Deutsche Forschungsgemeinschaft (DFG) grants EXC307 (CIN – Werner Reichardt Centre for Integrative Neuroscience), INST 37/967-1 FUGG, INST 37/1254-1 FUGG, and the Human Frontier Science Program (HFSP) Young Investigator grant RGY0079.

## Author contributions

Conceptualization: T.C.H., T.T. and A.B.A. Data Curation: T.C.H. Formal Analysis: T.C.H. Funding Acquisition: T.T. and A.B.A. Investigation: T.C.H., B.S., P.W. and A.W. Methodology: T.C.H., D.-S.B., Y.Z., T.T. and A.B.A. Project Administration: T.C.H. and A.B.A. Resources: T.C.H., D.-S.B. and Y.Z. Software: T.C.H and Y.Z. Supervision: A.B.A. Validation: T.C.H. Visualization: T.C.H. Writing – Original Draft Preparation: T.C.H. Writing – Review & Editing: T.C.H., D.-S.B. T.T. and A.B.A.

## Supplementary Information

**S-Table 1.**
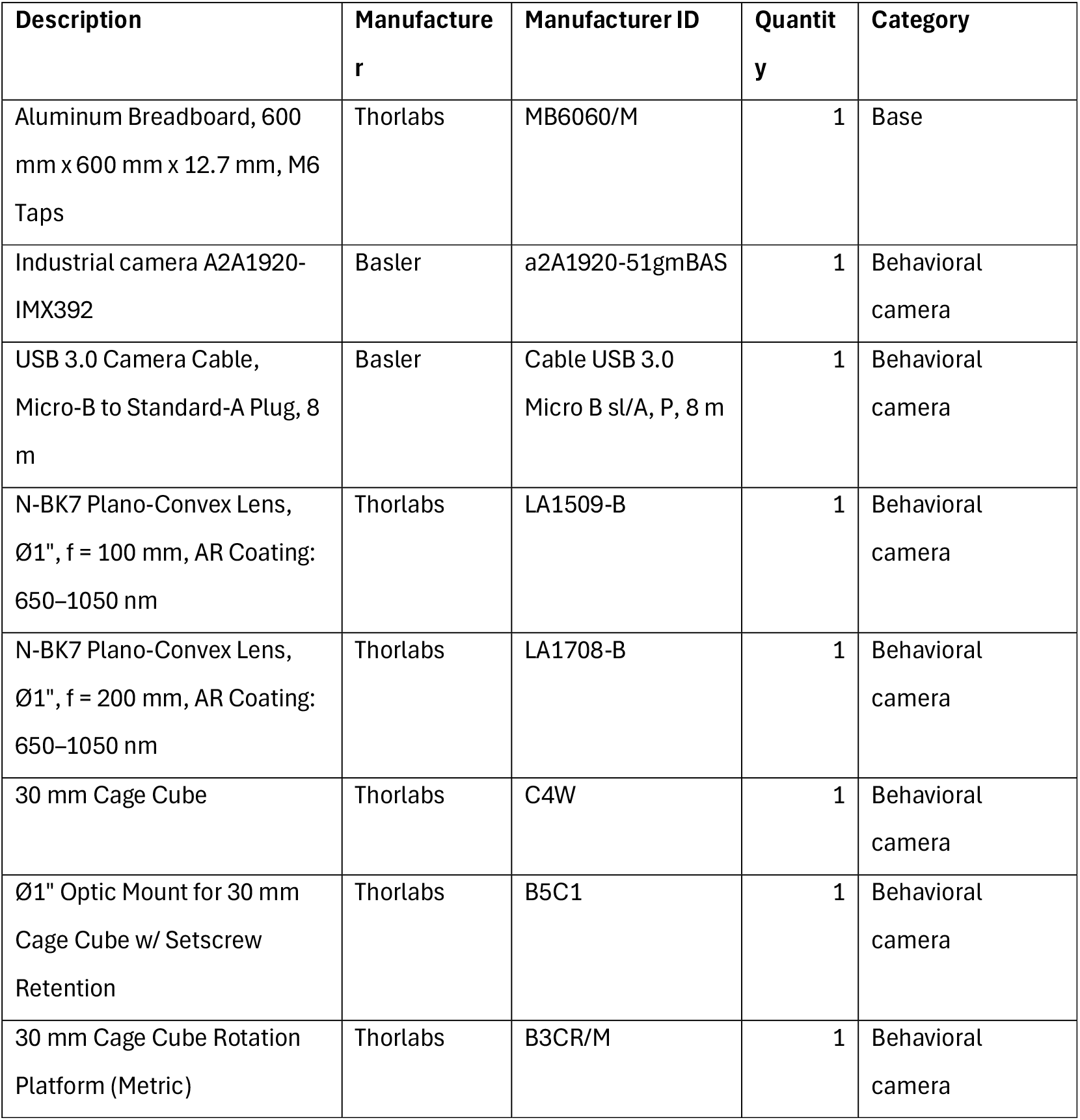
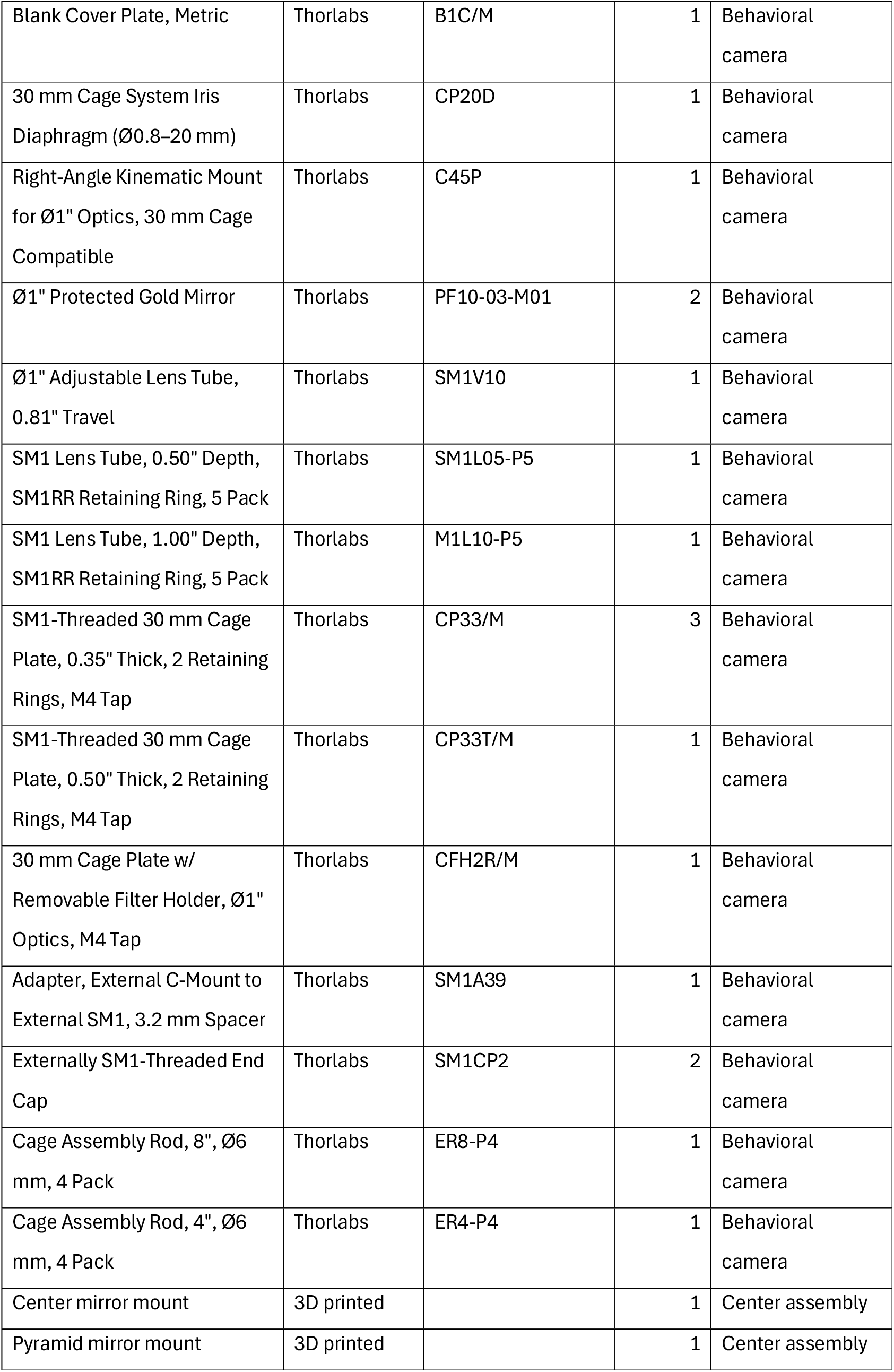
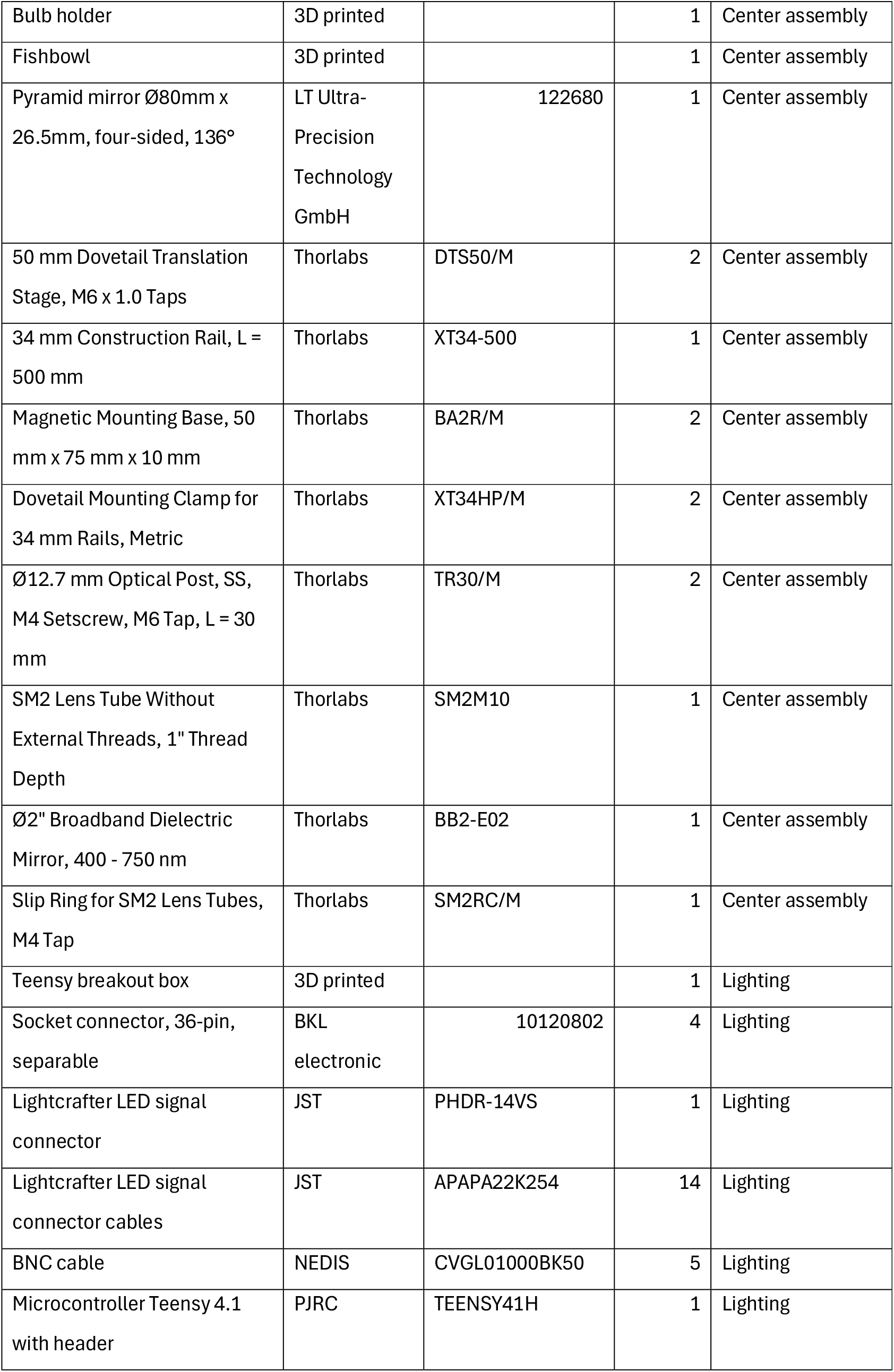
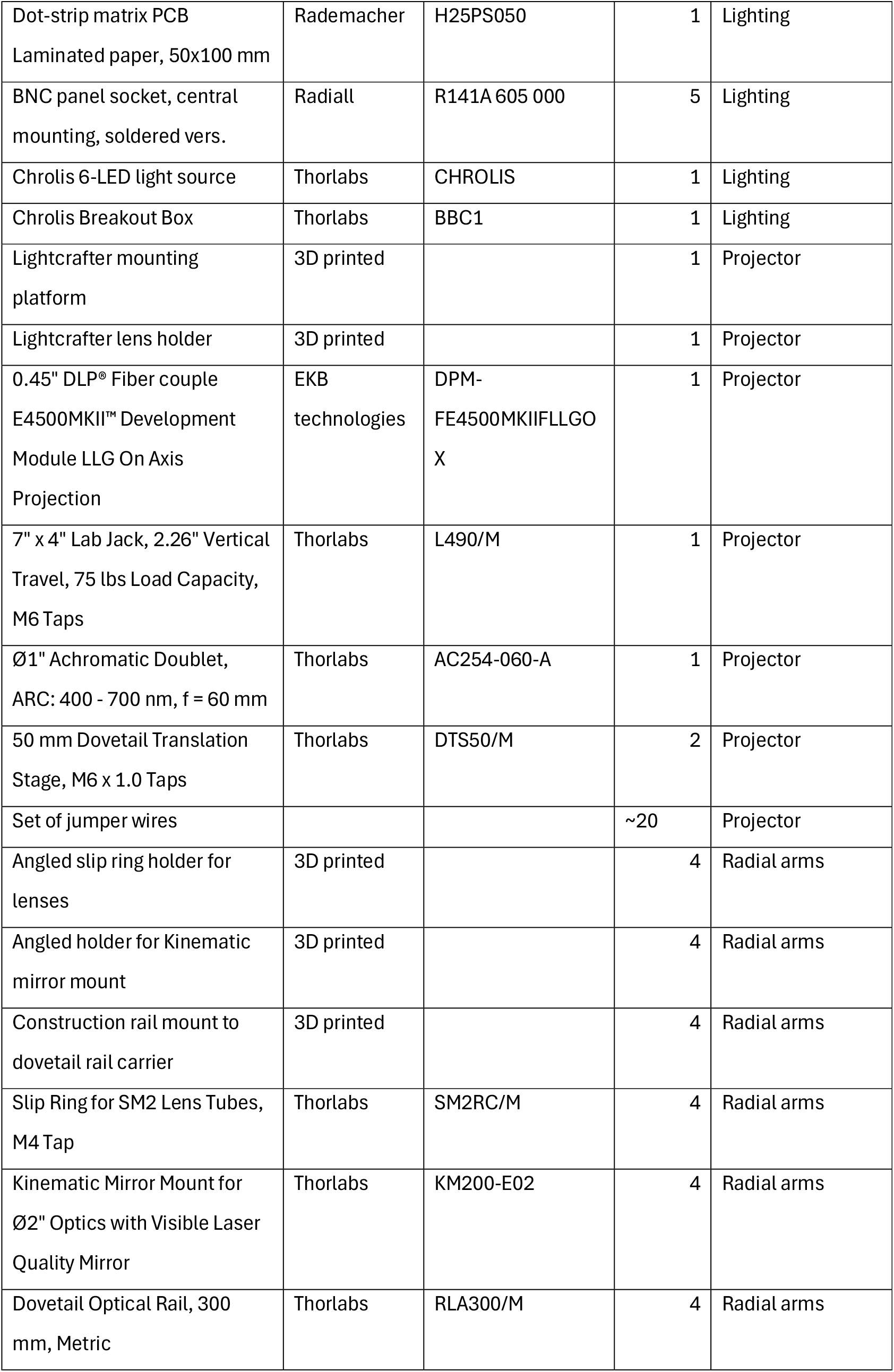
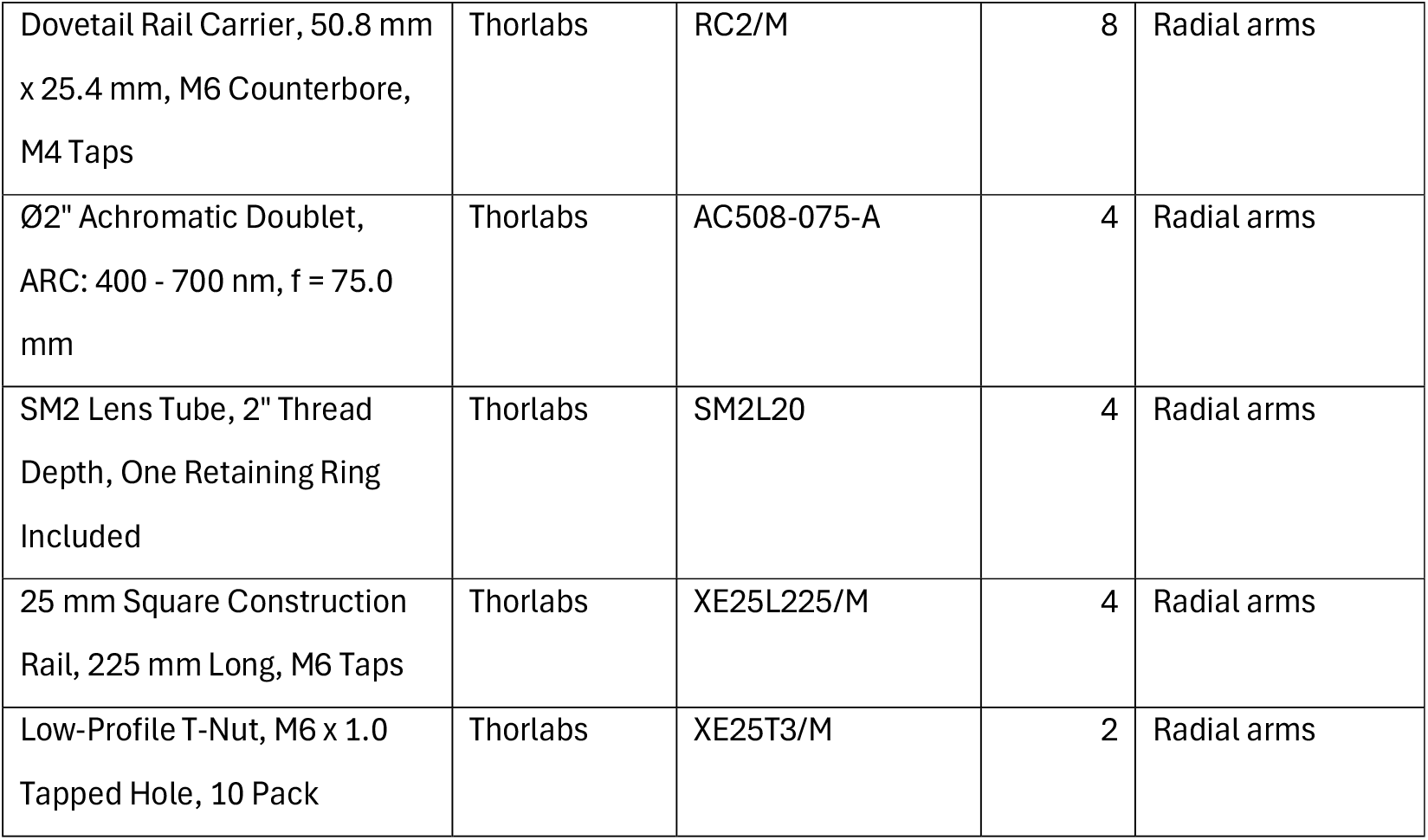
Bill of materials for stimulation setup.

**S-Table 2.** Irradiance measurements [μW/cm^2^] of the visual stimulation during projection of a 90° by 90° checkerboard at different zebrafish cone-optimized wavelengths of 625nm (red), 475m (green) and 420nm (blue). Measurements were taken without blanking of the stimulation LEDs and at an elevation of +22.5°. The measurements denoted with 0%(X) are measurements on the dark patches of the checkerboard for a given bright patch relative intensity of X (i.e. they show the background luminance on dark patches).

|  | **E<sub>max</sub>** |  |  |  | **E<sub>min</sub>** |  |  |  |
| --- | --- | --- | --- | --- | --- | --- | --- | --- |
|  | **100.00%** | **75.00%** | **50.00%** | **25.00%** | **0%(100%)** | **0%(75%)** | **0%(50%)** | **0%(25%)** |
| **Red** | 13.5 | 6.5 | 4.0 | 1.7 | 2.2 | 0.8 | 0.5 | 0.1 |
| **Green** | 6.4 | 4.0 | 2.4 | 1.0 | 0.6 | 0.4 | 0.1 | 0.0 |
| **Blue** | 4.1 | 2.2 | 1.3 | 0.3 | 0.3 | 0.0 | 0.0 | 0.0 |

**S-Table 3.**
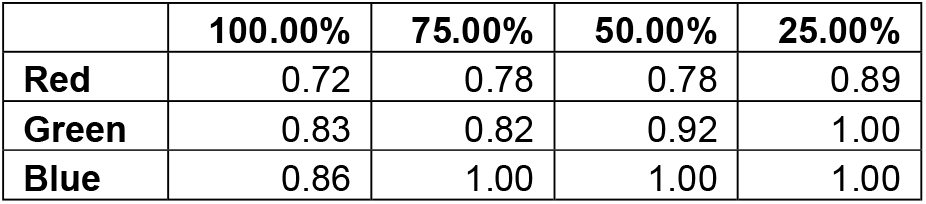
Michelson contrasts of the visual stimulation during projection of a 90° by 90° checkerboard at different zebrafish cone-optimized wavelengths of 625nm (red), 475m (green) and 420nm (blue).

|  | **100.00%** | **75.00%** | **50.00%** | **25.00%** |
| --- | --- | --- | --- | --- |
| **Red** | 0.72 | 0.78 | 0.78 | 0.89 |
| **Green** | 0.83 | 0.82 | 0.92 | 1.00 |
| **Blue** | 0.86 | 1.00 | 1.00 | 1.00 |

**S-Figure 1.**
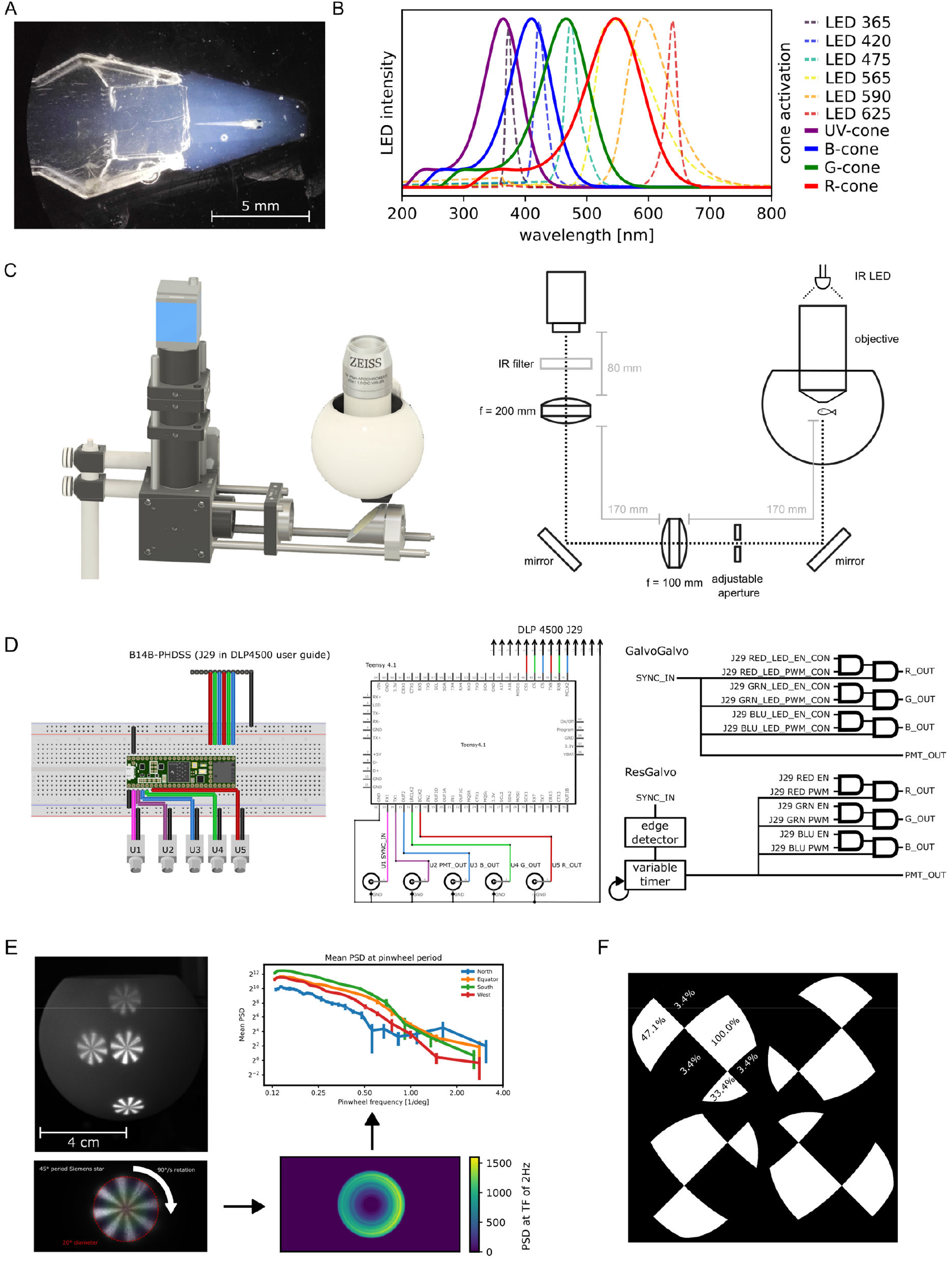
Overview of setup, microcontroller wiring and circuit logic. **A**: image of embedded zebrafish larva in low-melting agarose (1.5%) on top of a stage made from 2.5% agarose. **B**: Approximate activation spectra of zebrafish cones (solid lines) and stimulation LED spectra (dashed lines) after Franke et al. (2019). **C**: Image of behavioral camera path, including fishbowl (left). Schematic drawing with dimensions of camera path (right) **D**: Left - wiring between the Lightcrafter E4500MKII’s J29 connector, a Teensy 4.1 microcontroller and the input/output BNC connectors. Output connectors for the red, green and blue light sources (U5, U4 and U3 respectively), optional gating signal for the photomultiplier tubes (U2) and synchronization signal input (U1). Middle - minimal wiring diagram of the circuit in A. Right - Teensy 4.1 microcontroller’s software logic for switching output signals. Note that the GalvoGalvo variant is designed to use a y-scanline synchronization signal (∼1kHz) and SYNC_IN may be inverted. The ResGalvo variant’s synchronization is typically tied to the frame rate of the scanner at ∼30Hz and uses a customizable timer to switch output signals. **E**: Calculation of the optical spatial resolution of the stimulation arena for one of the four identical light paths. Calculation was done using Siemens stars at four different positions (north, equator, south and west) to compare the location-based uniformity of spatial resolution. Top right shoes a mean power spectral density low-pass filter function for each location on the sphere. **F**: Relative luminance power measurements for bright and dark subfields in a global checkerboard pattern (spatial period 90°) and for different positions on the clover projection pattern. Position-dependent luminance difference can be corrected by automatic stimulus preprocessing in the stimulation software (Hladnik, 2025)

**S-Figure 2.**
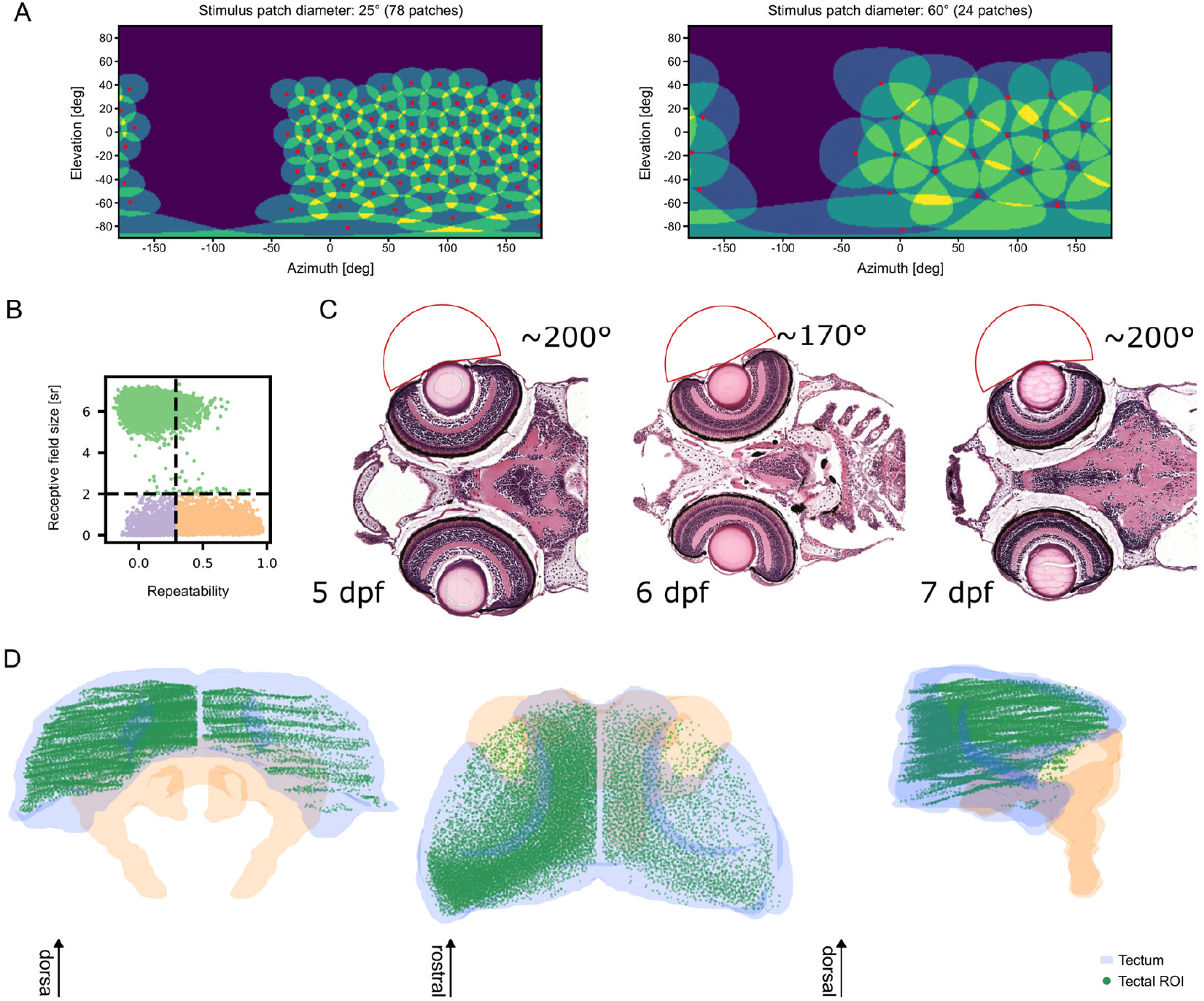
Supplementary information on spatial receptive field mapping experiments. **A**: Illustration of patch centers (red dots) and stimulation areas (yellow shaded area) that were used to present the moving foreground stimulus in front of the static background. Projection is a UV projection. **B**: Filtering parameters for selection of valid regions of interest for spatial receptive field analysis. Repeatability v. estimated receptive field size in steradians. Dashed lines show imposed threshold. Green shows all ROI’s excluded for their RF size; purple all that were excluded for their low trial-to-trial repeatability and orange are all cells included for further analysis. **C**: Illustration of approximate, maximally possible visual angles for which light rays might still be able to hit the eye’s lens and thus be refracted towards the retina. Image from PennState BioAtlas (Bio-Atlas -Bio-Atlas, n.d.; Copper et al., 2018). **D**: Anatomical overview of all ROI positions inside the tectum (green dots). Tectum (blue) and pretectum (orange) for reference.

**S-Figure 3:**
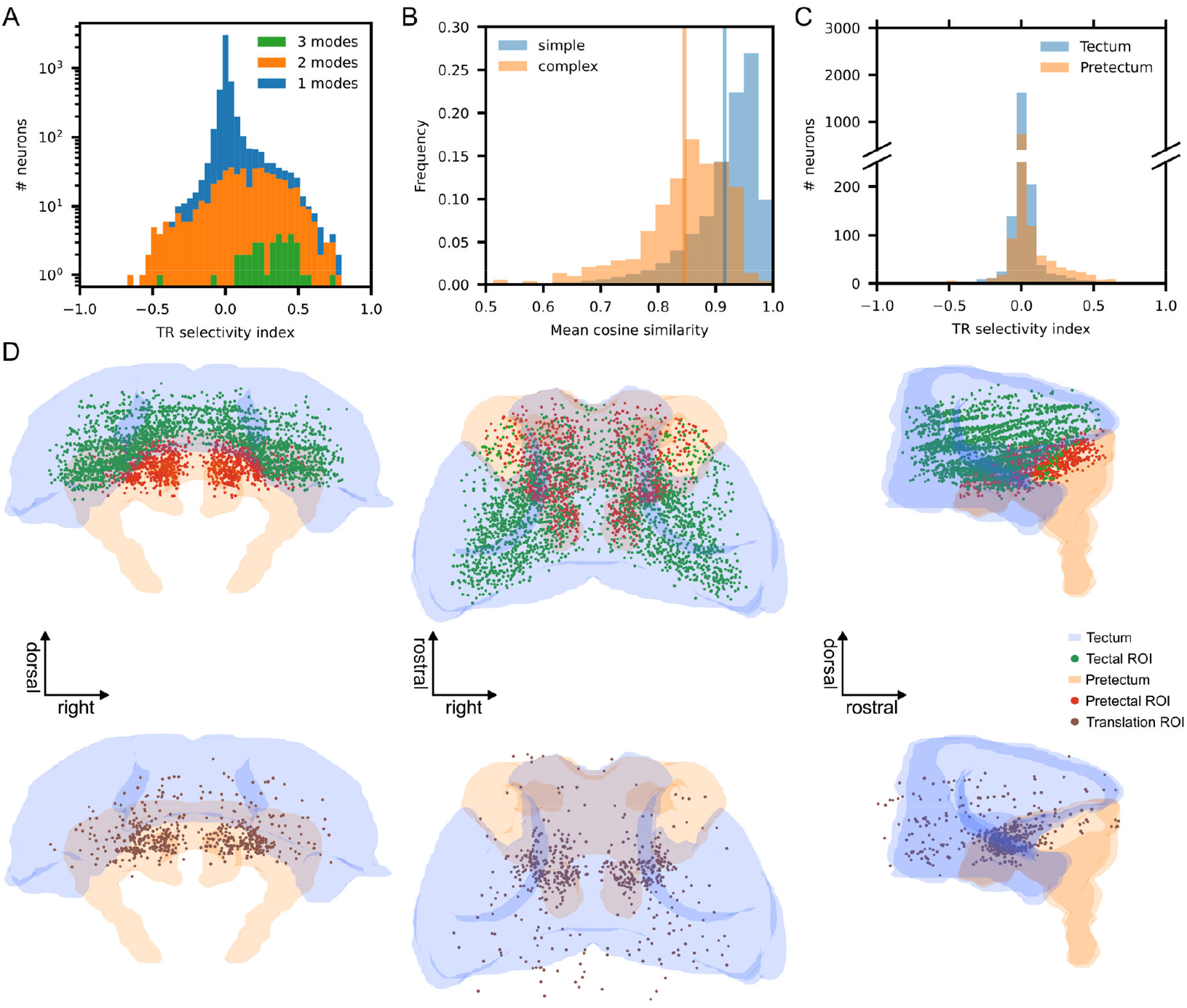
Supplementary information on CMN motion receptive field mapping experiment. **A**: Distribution of translation/rotation (TR) selectivity indices separated by the number of modes (subfields) of the respective cell’s receptive field. **B**: Frequency of cosine similarities based on their receptive field type. Simple cells (1 mode/subfield) had on average higher cosine similarities than complex cells (> 1 subfield). **C**: Distribution of translation/rotation (TR) selectivity indices for all cells, separated in color by anatomical location. **D**: Anatomical distribution of tectal, pretectal and translation-selective cells included in the analysis in Figure 3.

**S-Figure 4.**
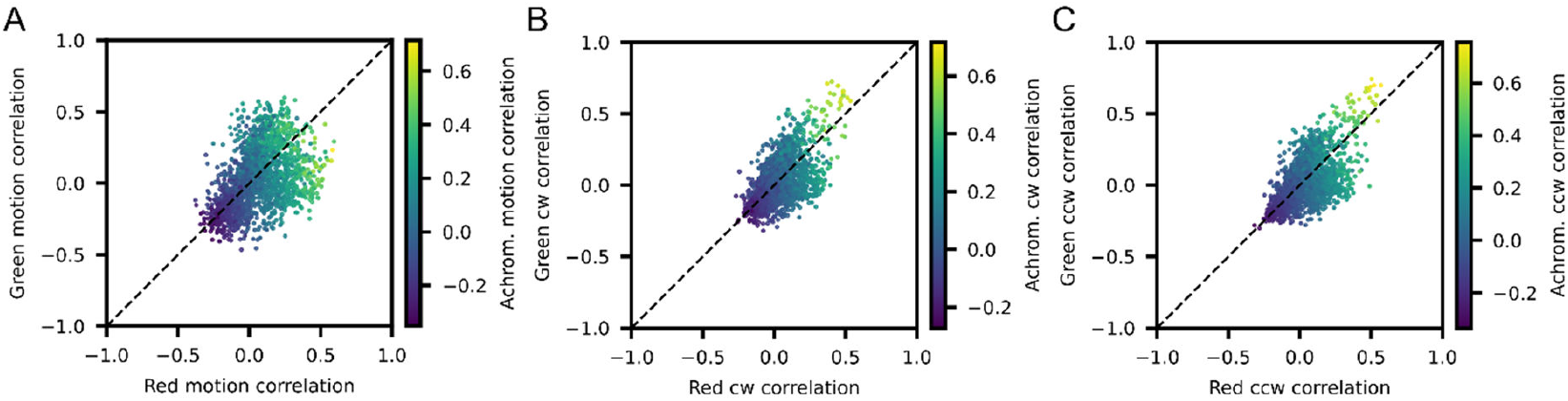
Supplementary information on global chromatic motion processing experiments. Correlations of all ROIs (n=1901) that showed reproducible responses across trial repeats (repeatability > 0.3) to achromatic contrast color coded as a function of red and green contrast correlations for non-directional motion (**A**), clockwise (**B**) and counterclockwise (**C**) motion types.

## Notes

https://github.com/thladnik/mariner-experimental-setup

### Competing Interest Statement

The authors have declared no competing interest.

