## Supplementary material for "MARINER: a surround visual stimulator for vision research in aquatic animals": MARINER light path schematic drawing

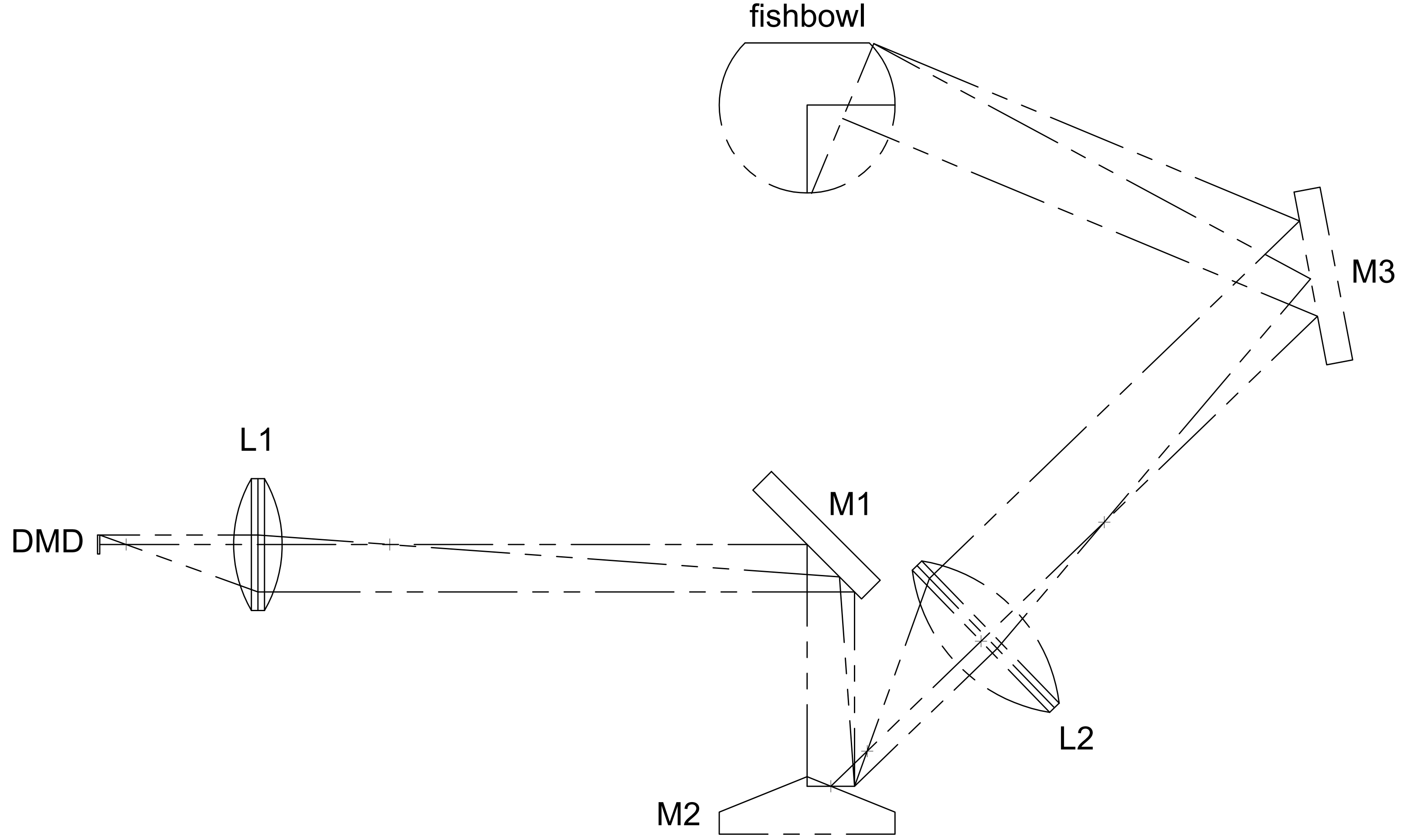


| Dept. | Technical reference | Created by<br>Tim Hladnik 2025-07-09 | Approved by |
|---|---|---|---|
| Unit mm | | Document type | Document status |
| | | Title<br>MARINER light path schematic | DWG No. |
| | | | Rev. \| Date of issue \| Sheet 1/4 |

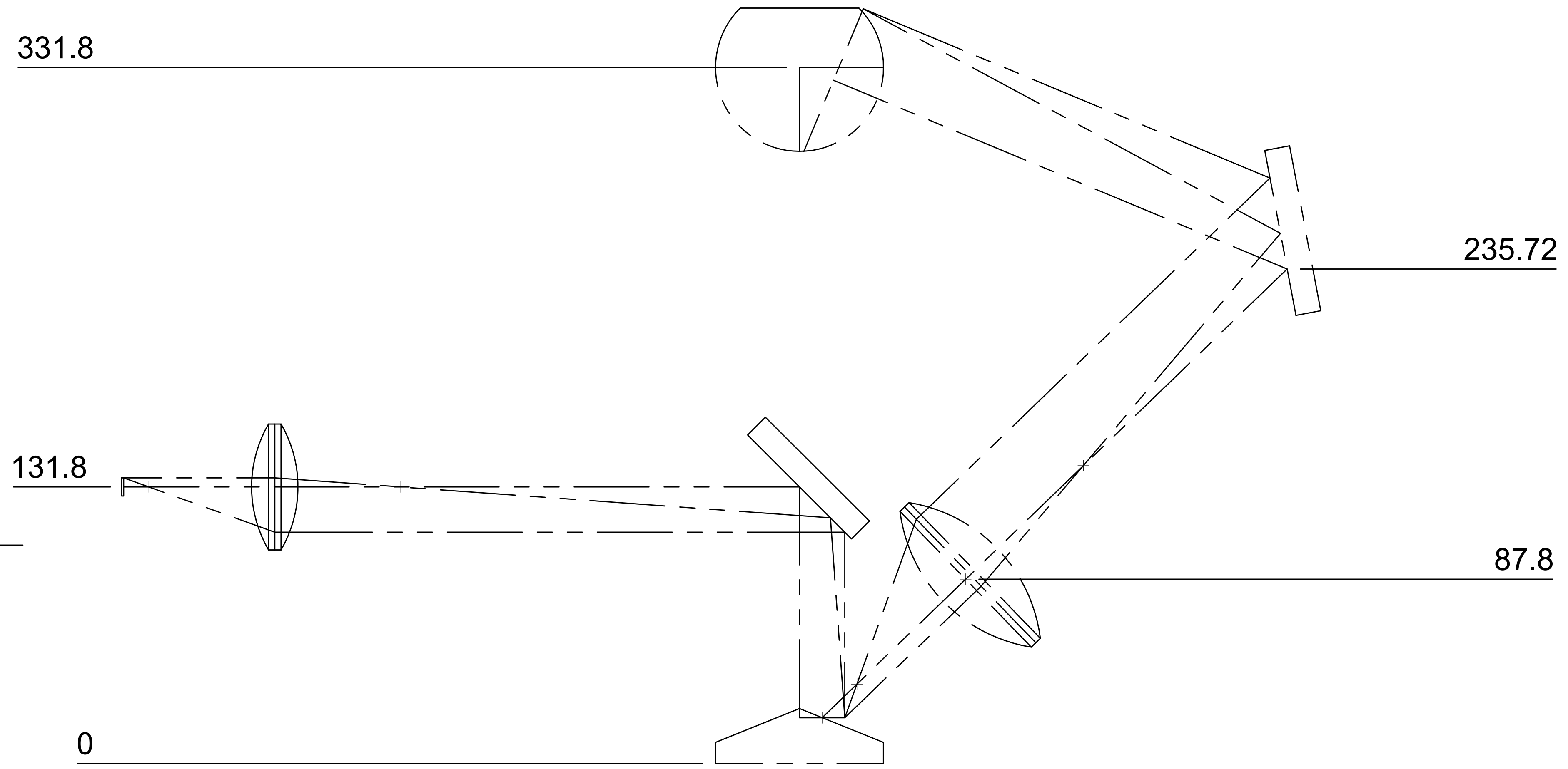


| Dept. | Technical reference | Created by | Approved by |
| --- | --- | --- | --- |
| | | Tim Hladnik 2025-07-09 | |
| Unit mm | | Document type | Document status |
| | | Title: MARINER light path schematic | DWG No. |
| | | | Rev. / Date of issue / Sheet 2/4 |

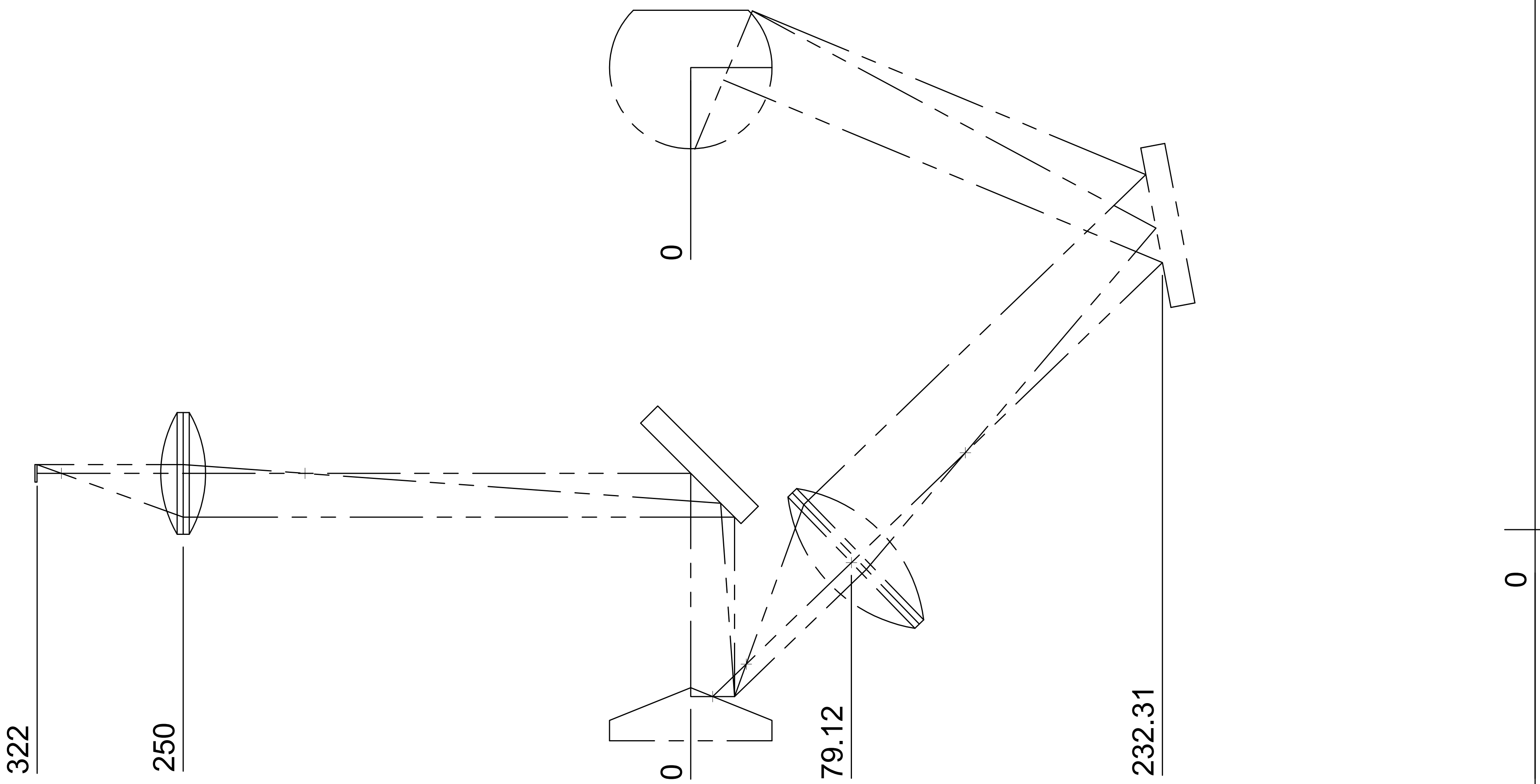


| Dept. | Technical reference | Created by Tim Hladnik 2025-07-09 | Approved by |
| --- | --- | --- | --- |
| Unit mm | | Document type | Document status |
| | | Title MARINER light path schematic | DWG No. |
| | | | Rev. / Date of issue / Sheet 3/4 |

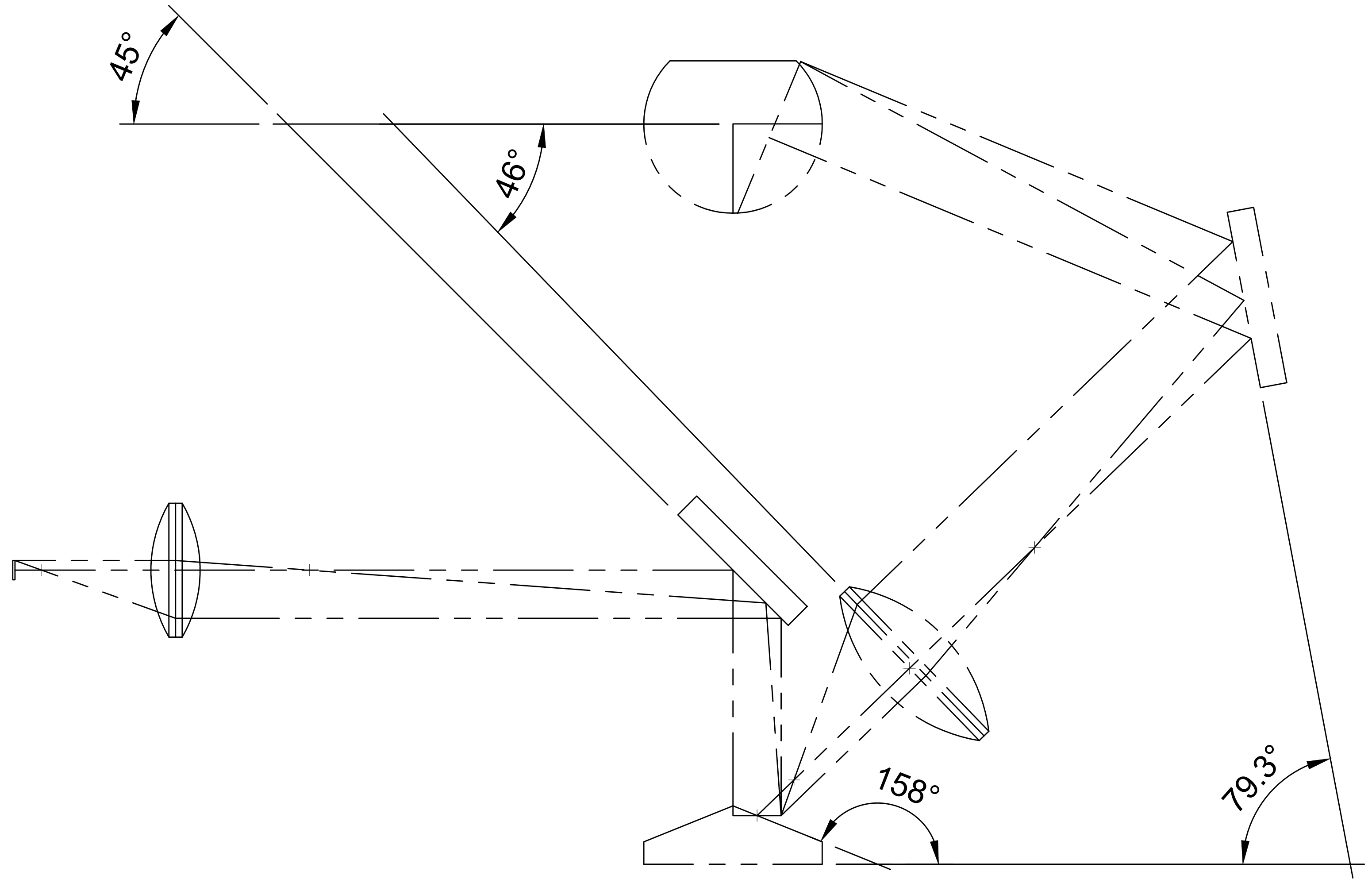


0

| | | | |
|---|---|---|---|
| Dept. | Technical reference | Created by<br>Tim Hladnik 2025-07-09 | Approved by |
| Unit mm | | Document type | Document status |
| | | Title<br>MARINER light path schematic | DWG No. |
| | | | Rev. \| Date of issue \| Sheet 4/4 |
